# Hydrophobic mismatch is a key factor in protein transport on the Tat pathway

**DOI:** 10.1101/2021.05.26.445883

**Authors:** Binhan Hao, Wenjie Zhou, Steven M. Theg

## Abstract

The twin-arginine translocation (Tat) pathway transports folded proteins across membranes in bacteria, thylakoid, plant mitochondria, and archaea. In most species, the active Tat machinery consists of three independent subunits, TatA, TatB and TatC. TatA and TatB from all bacterial species possess short transmembrane alpha-helices (TMHs), both of which are only fifteen residues long in *E. coli*. Such short TMHs cause a hydrophobic mismatch between Tat subunits and the membrane bilayer. Here, by modifying the length of the TMHs of *E. coli* TatA and TatB, we assess the functional importance of the hydrophobic mismatch in the Tat transport mechanism. Surprisingly, both TatA and TatB with as few as 11 residues in their respective TMHs are still able to insert into the membrane bilayer, albeit with a decline in membrane integrity. Three different assays, both qualitative and quantitative, were conducted to evaluate the Tat activity of the TMH length mutants. Our experiments indicate that the TMHs of TatA and TatB appear to be evolutionarily tuned to 15 amino acids, with activity dropping off with any modification of this length. We believe our study supports a model of Tat transport utilizing localized toroidal pores that form when the membrane bilayer is thinned to a critical threshold. In this context, the 15-residue length of the TatA and TatB TMHs can be seen as a compromise between the need for some hydrophobic mismatch to allow the membrane to reversibly reach the threshold thinness required for toroidal pore formation, and the permanently destabilizing effect of placing even shorter helices into these energy-transducing membranes.

## Introduction

The twin-arginine translocation (Tat) pathway, which is found in prokaryotes, archaebacteria, chloroplasts thylakoids and some mitochondria, is able to transport multiple substrate proteins across the membrane lipid bilayers. In chloroplasts, this pathway is responsible for the transport of a number of essential proteins, including two of the three subunits of the oxygen-evolving complex (Clark and Theg, 1997). In bacteria, the Tat pathway serves several critical biological processes, including electron transport, cell division, cell wall formation, stress tolerance, and pathogenesis (Ize et al., 2003; Palmer et al., 2005). The Tat pathway has the following unusual characteristics. First, it has the ability to transport folded proteins, which is fundamentally different from, for instance, the mitochondrial import and the ubiquitous Sec pathways. Second, Tat pathway substrates have a unique cleavable signal peptide which carries a nearly invariant pair of arginines (-R-R-) (New et al., 2018). Third, this pathway uses the protonmotive force (PMF) as the sole energy source, with no contribution from NTP hydrolysis (Braun et al., 2007). Fourth, the Tat pathway acts in an ion-tight manner while transporting substrates of different sizes (Asher and Theg, 2021; Teter and Theg, 1998). Fifth, the complete translocation machinery assembles only transiently during the transport event (Mori and Cline, 2002). Even though Tat pathway can transport folded proteins with different sizes, the Tat translocon, in most species, involves only three functionally independent subunits, TatA, TatB and TatC. It has been shown that TatA, TatB and TatC form a protein complex which serves as the receptor for Tat signal peptide (Gérard and Cline, 2007; Johann et al., 2017; Taubert et al., 2015). The assembly into a functional translocon that includes TatA depends on substrate binding and the PMF. Finally, even though TatB and TatC are present in a 1:1 stoichiometry, TatA joins the complex in variable stoichiometries (Leake et al., 2008).

The structures of the three Tat subunits have been reported (Hu et al., 2010; Pettersson et al., 2018; Rollauer et al., 2012; Zhang et al., 2014b). TatA and TatB share an overall “L-shape” conformation composed of an N-terminal undefined short region located in the periplasm in bacteria, a remarkably short transmembrane alpha helix (TMH), a hinge region followed by one or more amphipathic helices (APH), and an unstructured C-terminal tail. In contrast, TatC has six transmembrane helices and is configured in a cupped hand shape with both N-terminal and C-terminal located in the cytoplasm.

Three models have been proposed for the mechanism of protein transport on the Tat pathway. In the first model, TatA is proposed to form a form-fitting proteinaceous pore around the incoming transport substrate (Frain et al., 2019; Gohlke et al., 2005). In a second model, TatA is thought to act as a co-enzyme that accumulates to activate a translocon built from a complex consisting of TatB and TatC (Hauer et al., 2017, 2013). A third model posits that under the influence of all Tat subunits, the substrate itself and the protonmotive driving force, the membrane thins locally until the bilayer breaks down with the formation of lipid-lined toroidal pores through which substrates traverse the membrane (Asher and Theg, 2021; Brüser and Sanders, 2003; Hou et al., 2018). Unlike proteinaceous pores in which amphipathic helices form water-filled channels (Berg et al., 2004; Tsirigotaki et al., 2017), in toroidal pores the lipid molecules wrap across the membrane bilayer to form channels toroidal in shape. A similar lipid toroidal pore model is thought to govern the translocation of antimicrobial peptides across biological membranes (Chen et al., 2003; Huan et al., 2020; Lee et al., 2013). It has been shown that toroidal pores are quite general since they can be induced not only by peptides (Brown and Hancock, 2006) but also by other chemicals (Gurtovenko and Anwar, 2007). The first and third models are minimally depicted in Figure 1; more detailed representations of toroidal pores and their role in membrane transport can be found in (Asher and Theg, 2021; Huan et al., 2020; Sengupta et al., 2008).

**Figure 1.**
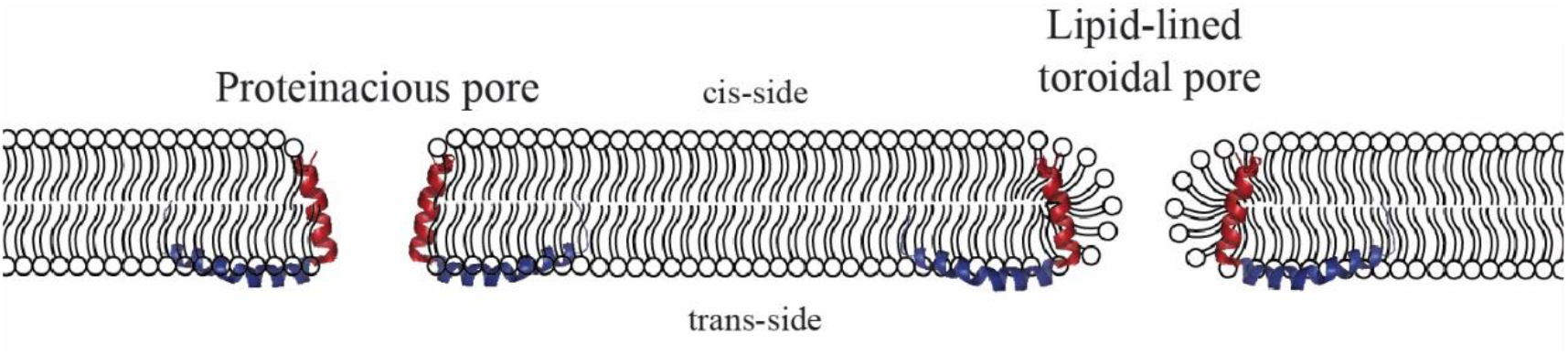
Two models of Tat protein transport. The L-shaped peptide represents the TatA TMH (red) and APH (blue); the TatA C-terminus is not drawn for simplicity. Left: a proteinaceous pore delineated by the TatA TMH (model 1 in text); right: a toroidal pore with TatA TMHs buried under the lipid head groups (model 3 in text).

Although there is no detailed information about the structure of the active Tat machinery, a special structural feature in the TatA and TatB TMHs potentially point to an active role of the membrane biophysics in the mechanism of Tat pathway. In *E. coli*, the TMHs of TatA and TatB only have 15 amino acids, respectively, which makes the length of TMHs (∼22.5 Å) much shorter than the normal thickness of the hydrophobic core of the membrane (30Å) (Mitra et al., 2004). The difference between the length of the TMHs and the thickness of membrane bilayer cause a hydrophobic mismatch effect. Many studies have shown that the activity of membrane proteins can be sensitive to such mismatch (Brandizzi et al., 2002; Milovanovic et al., 2015; Parton et al., 2011). The possible consequences of the hydrophobic mismatch between short TMHs and lipid bilayers are various and depend on the overall topology of the proteins. One of the outcomes is protein aggregation or oligomerization (Killian, 1998), which can cause proximal thinning of the membrane bilayer. Such protein oligomerization phenomenon is also observed in TatA, which forms higher order structures in the resting state of Tat transport (Dabney-Smith et al., 2006; Palmer and Berks, 2012). It remains unclear that whether such hydrophobic mismatch between Tat subunits and membrane bilayer is necessary for the Tat pathway. A previous study showed that the TMH of TatA (without the APH) can destabilize the membrane in inverted membrane vesicles (IMVs), even though the full-length TatA does not show a similar proton leakage effect. Such membrane destabilization could potentially be involved in the formation of transport-competent toroidal pores (Hou et al., 2018).

In the present study, we investigate the hydrophobic mismatch between Tat subunits and the membrane bilayer by modifying the length of the TMHs of *E. coli* TatA and TatB. Up to five amino acids were added to the TMHs at three different loci to decrease the hydrophobic mismatch. Conversely, up to four amino acids were deleted from the TMHs to increase the hydrophobic mismatch. The effects of these changes in TMH lengths were examined by three different measures of Tat activity, both qualitative and quantitative. We found that the hydrophobic mismatch between Tat subunits and membrane bilayer appears to be optimized for maximal Tat activity. We further found that decreasing the length of the TatA TMH caused leakage of protons, and presumably other ions, across the membrane. These findings offer the insights into functional importance of the unusually short TMHs of TatA and TatB for the mechanism of Tat translocation.

## Results

### A conserved 12 amino acid-long hydrophobic region is present in TatA and TatB across different species

The *E. coli* TatA and TatB TMHs each consist of 15 residues (Ile6-Phe20 in the TatA and Phe6-Leu20 in the TatB) according to the NMR structures (Zhang et al., 2014a, 2014b). In *Bacillus subtilis*, a gram-positive bacteria, the TatAd also includes a TMH with 15-residues (Ile7-Phe21) (Hu et al., 2010). To assess whether the length of these short TMHs is conserved across different species, 122 TatA and 60 TatB sequences from bacteria, chloroplasts and mitochondria were aligned by MUSCLE (Figure 2A and 2B). Consistent with the previous literature (Barrett et al., 2003), Phe(F)-Gly(G) and Gly(G)-Pro(P) motifs were observed in the TatA and TatB alignments, respectively, and a conserved polar amino acid locus (#8 in the TatA and TatB sequence logo plots) was also observed. However, the precise TMH length of TatA/B sequences could not be determined due to lack of structural information for each Tat protein. Furthermore, since there are conserved hydrophilic amino acids present at 8^th^ position in TatA/B (*E. coli* TatA/B numbering, Figure 2A and 2B), it is also difficult to estimate the TMH length. Even though it is hard to predict accurate length for each TatA/B sequence, a 12-residue hydrophobic region between the polar residue (position #8) and the glycine (position #21) is found to be extremely conserved among all TatA and TatB sequences analyzed. Such conservation of the length of the TatA and TatB TMHs across species which display somewhat different membrane bilayer thicknesses (Mitra et al., 2004; Perkins et al., 1997; Pribil et al., 2014) suggests that the length of the TatA and TatB TMH has potential significance for the Tat transport mechanism.

**Figure 2.**
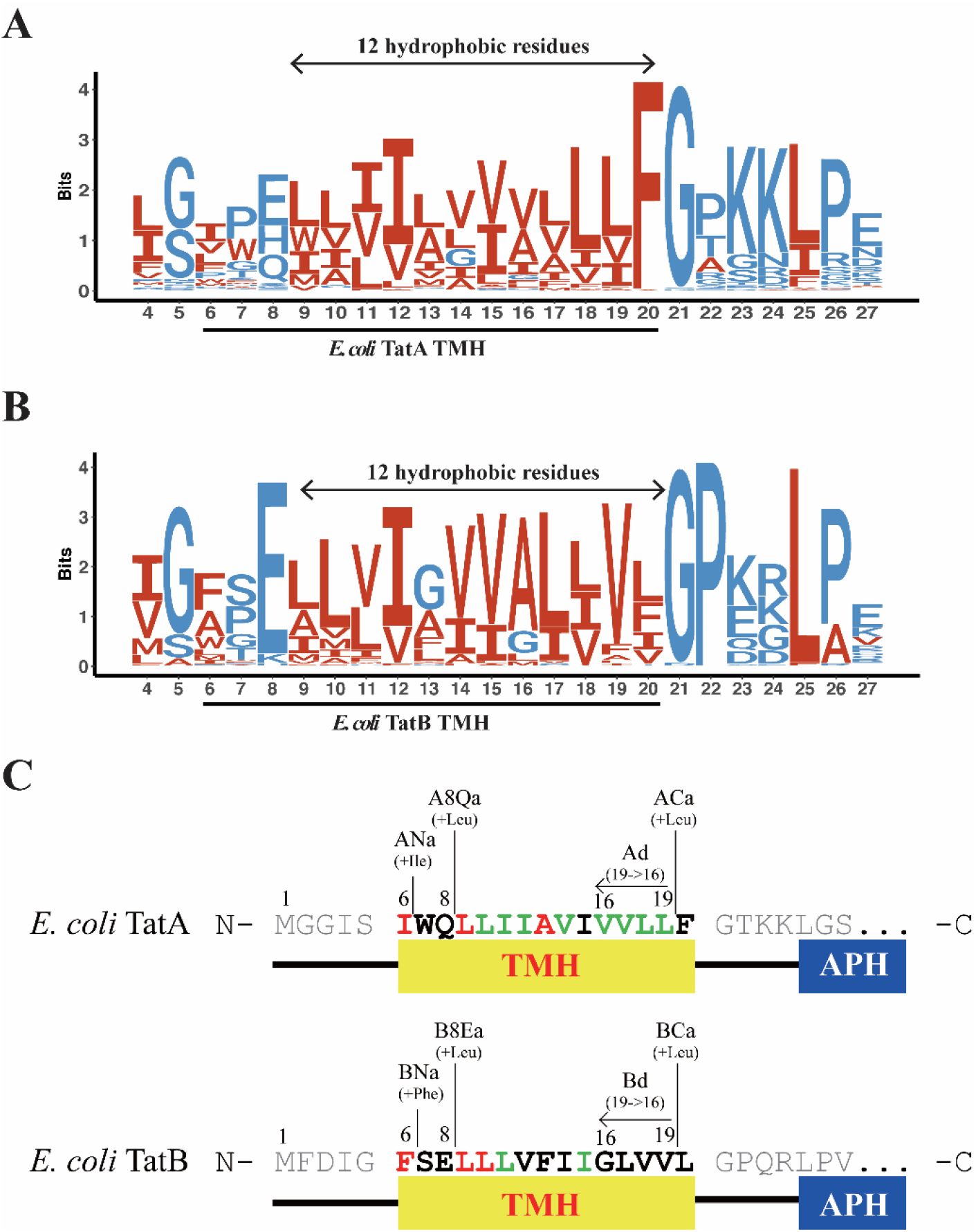
Sequence alignments of TatA and TatB TMH and schematic diagrams for mutant design. (**A,B**) Sequence logos for prokaryotic TatA and TatB alignments, respectively. 122 TatA sequences and 60 TatB sequences were downloaded from GenBank and subjected to multiple sequence alignment using MUSCLE. Sequence logos were subsequently generated using ggseqlogo in RStudio, where *E. coli* TatA and TatB numbering was used to denote residue locations. Hydrophobic residues were represented in red, and hydrophilic residues were represented in blue. An invariant 12-residue long hydrophobic region is present in both TatA and TatB and is highlighted by arrows. (**C**) Schematic diagram for the design and naming of *E. coli* TatA and TatB TMH mutants. In prose, the first letter in the mutant name, A or B, refers to TatA or Tat B, respectively. The next letters and numbers refer to the position of the change; N means at the TMH N-terminus, 8Q or 8E means after the polar amino acid at position 8, C means at the TMH C-terminus; a or d means addition or deletion, respectively, with all deletions occurring at the TMH C-termini. The final number refers the number of amino acids added or deleted at the specified position. Thus, ANa3 is the mutant construct wherein <u>3</u> amino acids were <u>a</u>dded at the <u>N</u>-terminus of the <u>TatA</u> TMH, A8Q2 is the construct wherein <u>2</u> amino acids were <u>a</u>dded after <u>Q8</u> in <u>TatA</u>, and Bd2 refers to the construct wherein <u>2</u> amino acids were <u>d</u>eleted from the C-terminus of the <u>TatB</u> TMH. The exact sequences of the different TMH mutants are shown below in Table I.

### TMHs with only 15 residues are not common

The 15-residue length of the TatA and TatB TMHs is remarkable in that they are expected to be longer (Saidijam et al., 2018) to span the 30 Å hydrophobic core of a typical membrane (Figure 3A). In order to understand how common such short TMH lengths are, we analyzed the TMH lengths in thousands of single-pass proteins from bacteria and chloroplasts. Figure 3B demonstrates that, as expected, such short-length TMHs are relatively rare, suggesting again that there is some functional significance to this feature of TatA and TatB.

**Figure 3.**
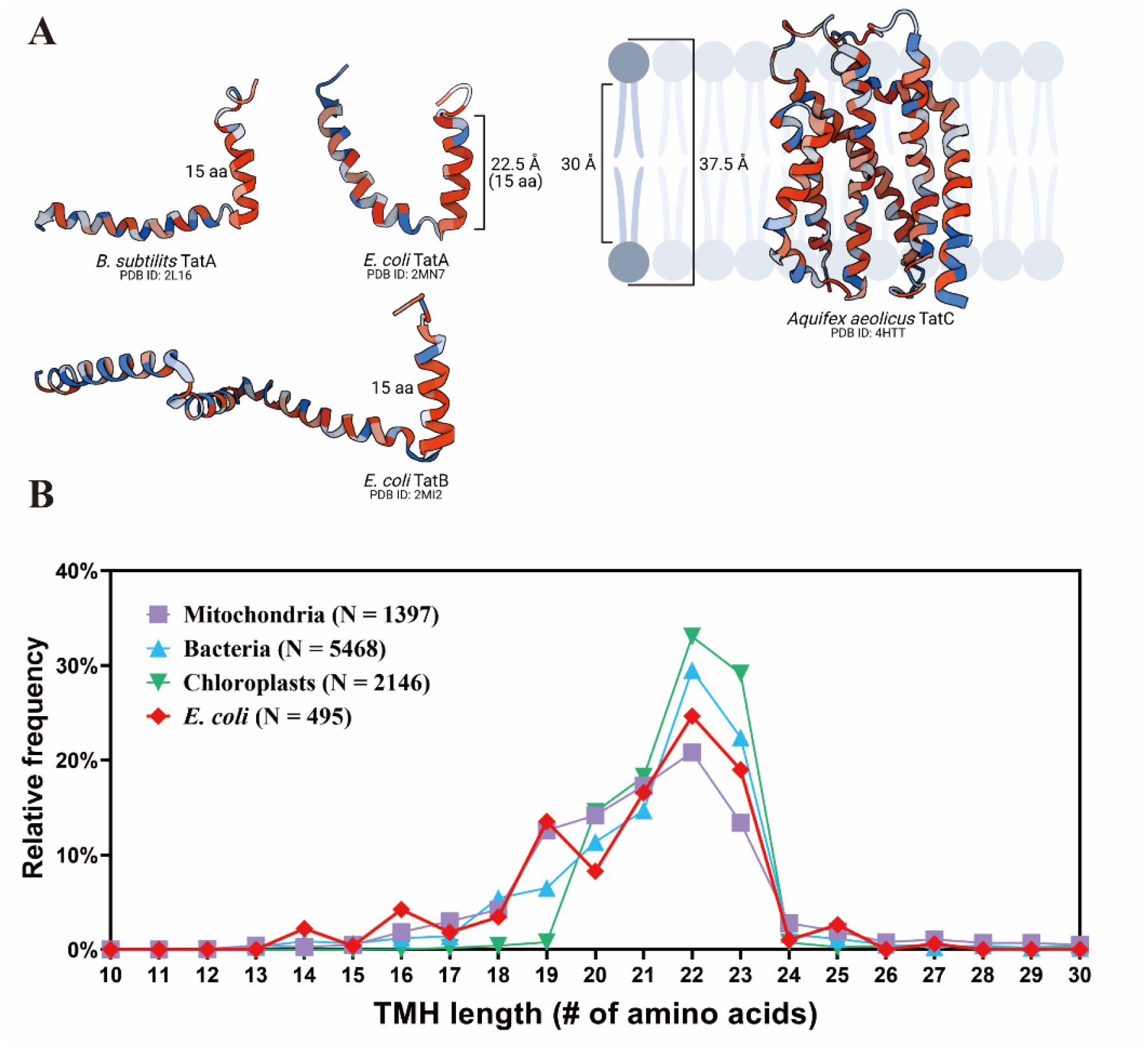
Frequency of short TMHs in selected organelles and organisms. (**A**) Representation of the hydrophobic mismatch between Tat subunits and the membrane bilayer. Protein structure was obtained from the Protein Data Bank. (**B**) Statistical analysis of the TMH length of proteins across different species. TMH length for each protein was predicted by TMHMM Server, v.2.0, which was then rounded to the nearest integer. Relative frequency in percentages was obtained by calculating the ratio of the number of proteins with predicted TMH at the indicated length to the total number of proteins in the corresponding category (mitochondria, bacteria, chloroplast, and *E. coli*). N, the total number of entries in each category. Further details are in Materials and Methods.

### Experimental modification and nomenclature for TMH-length modifications in TatA and TatB

To better understand how subunit hydrophobic mismatch contributes to Tat transport, we modified the lengths of the TMHs by adding or deleting amino acids from *E. coli* TatA and TatB. Four structural and functional principles were considered to minimize the effect on the overall topology of the TatA and TatB subunits when lengthening or shortening the TMHs. First, as the TMHs are α−helices, modifying the number of amino acids contained therein changes not only the length but also the potential protein-interacting helix faces. In order to minimize the relative rotation from subunit interfaces and APH orientation, amino acids were added close to the helix termini, rather than in the middle. Second, for the same reason, amino acid deletions were performed at the helix C-terminus. Third, we avoided deleting the conserved residues and functional groups in the TMHs. Fourth, we added the same amino acids as the one adjacent to the addition location. Based on those principles, three different loci were selected for the addition of one to five amino acids to lengthen the TMHs, and one location was selected to delete one to four amino acids to shorten the TMHs (Figure 2C). The various length mutants include the following: First, the Tat<u>A N</u>-terminus <u>a</u>ddition (**ANa**) group and the Tat<u>B N</u>-terminus <u>a</u>ddition (**BNa**) group in which amino acids were added at the extreme N-terminus of the TMHs. Second, the Tat<u>A</u> 8^th^ <u>Glutamine a</u>ddition (**A8Qa**) group and the Tat<u>B</u> 8^th^ <u>Glutamate a</u>ddition (**B8Ea**) group in which residues were added immediately following the polar amino acid in the TatA or TatB TMHs. Third, the Tat<u>A C</u>-terminus <u>a</u>ddition (**ACa**) group and the Tat<u>B C</u>-terminus <u>a</u>ddition (**BCa**) group, in which residues were added at the extreme C-terminus of the TMHs before the conserved 19^th^ Phe in TatA or the 19^th^ Leu, in the TatB. For deletion mutants, up to four amino acids before the 19^th^ Phe (in TatA) or the 19^th^ Leu (in TatB) were deleted step by step from the C- to N-terminus direction and are named the Tat<u>A d</u>eletion (**Ad**) group and Tat<u>B d</u>eletion (**Bd**) group. For example, “Ad2” represents the mutant whose 19^th^ Valine and 18^th^ Valine from the TatA TMH were deleted. In this study, we used the low copy pTat101 (Kneuper et al., 2012) as the parental plasmid, which constitutively expresses TatA, TatB and TatC. This plasmid was chosen to express Tat subunits at native expression levels and avoid the potential phenotype variation due to overexpression of Tat subunits. Table 1 shows the detailed design described above for all the mutants.

**Table 1.**
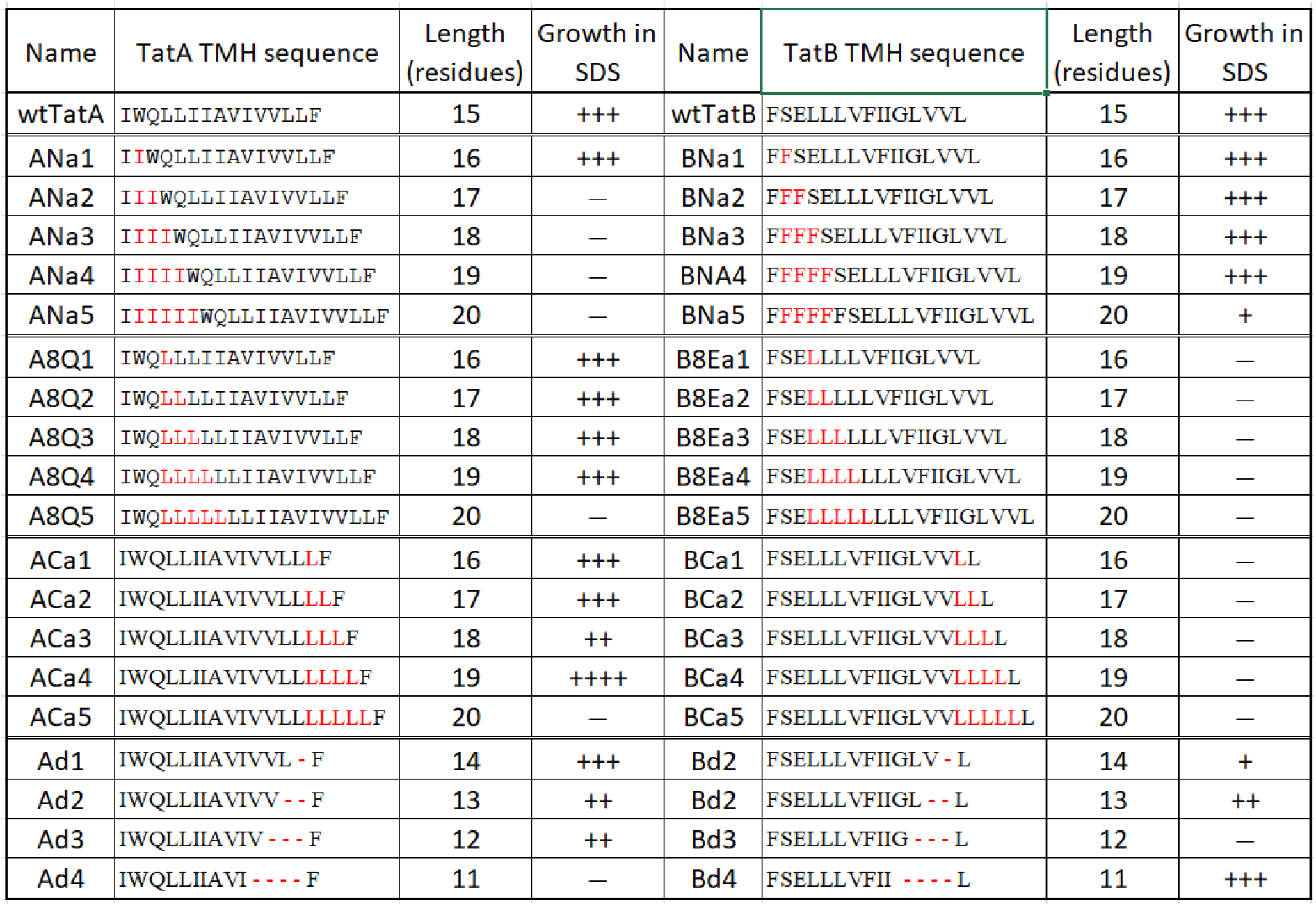
Summary of growth performance in the presence of SDS for TatA and TatB mutants. Growth rformance is described by the following symbols. +++, cells exhibited higher than 50% survival rcentage in 10% SDS; ++, cells exhibited higher than 50% survival percentage in 5% SDS; +, cells hibited higher than 10% survival percentage in 5% SDS; –, cells exhibited less than 10% survival rcentage in 5% SDS. Survival ratios were calculated as OD at 600 nm of cells grown for five hours in the esence of SDS to those grown in the absence of SDS. Detailed survival rates in media with different cocentrations of SDS are shown in Supplemental Figure 2. The sequences show the positions and amino acids added (red letters) or deleted (red —) for the various mutants.

### TatA and TatB deletion mutants exhibited lower membrane insertion stability

An obvious challenge for membrane proteins with short TMHs is correct and stable insertion into the membrane. To assess the membrane stability of our mutant proteins, membranes were isolated from whole cells and treated with 100 mM sodium carbonate to wash off non-integrated membrane proteins (Kim et al., 2015). According to the Western-blot results (Supplemental Figure 1), all the TatA and TatB addition mutants exhibited the expected stable membrane insertion ability. In contrast, a decrease in the membrane abundance was observed in the Ad and Bd deletion groups (Figure 4A and 4B). The amount of membrane-embedded TatA in the Ad group averaged approximately 20% of the amount found in the wild-type (Figure 4A). Similarly, all Bd mutants displayed less abundance compared to the wild-type TatB (Figure 4B). It is surprising that the Ad4 TatA and Bd4 TatB, which have only 11 amino acids in the TMH, were still detected in the membrane fraction. Previous research (Behrendt and Brüser, 2014) showed that TatB and TatC tend to form a TatBC complex in a one to one stoichiometry in the resting state of the Tat translocon. To test whether the TatC contributes to the membrane stability of the TatB deletion group, TatC was knocked out in the Bd4 mutant (i.e., Bd4*ΔtatC*). Even though a significant amount of Bd4 TatB was present in the membrane fraction when TatC was present, no Bd4 TatB was detected in the membrane of the Bd4*ΔtatC* mutant (Figure 4C). This is clear evidence that TatC stabilizes TatB in the membrane such that TatB can be embedded when its TMH is too short to remain in the membrane on its own.

**Figure 4.**
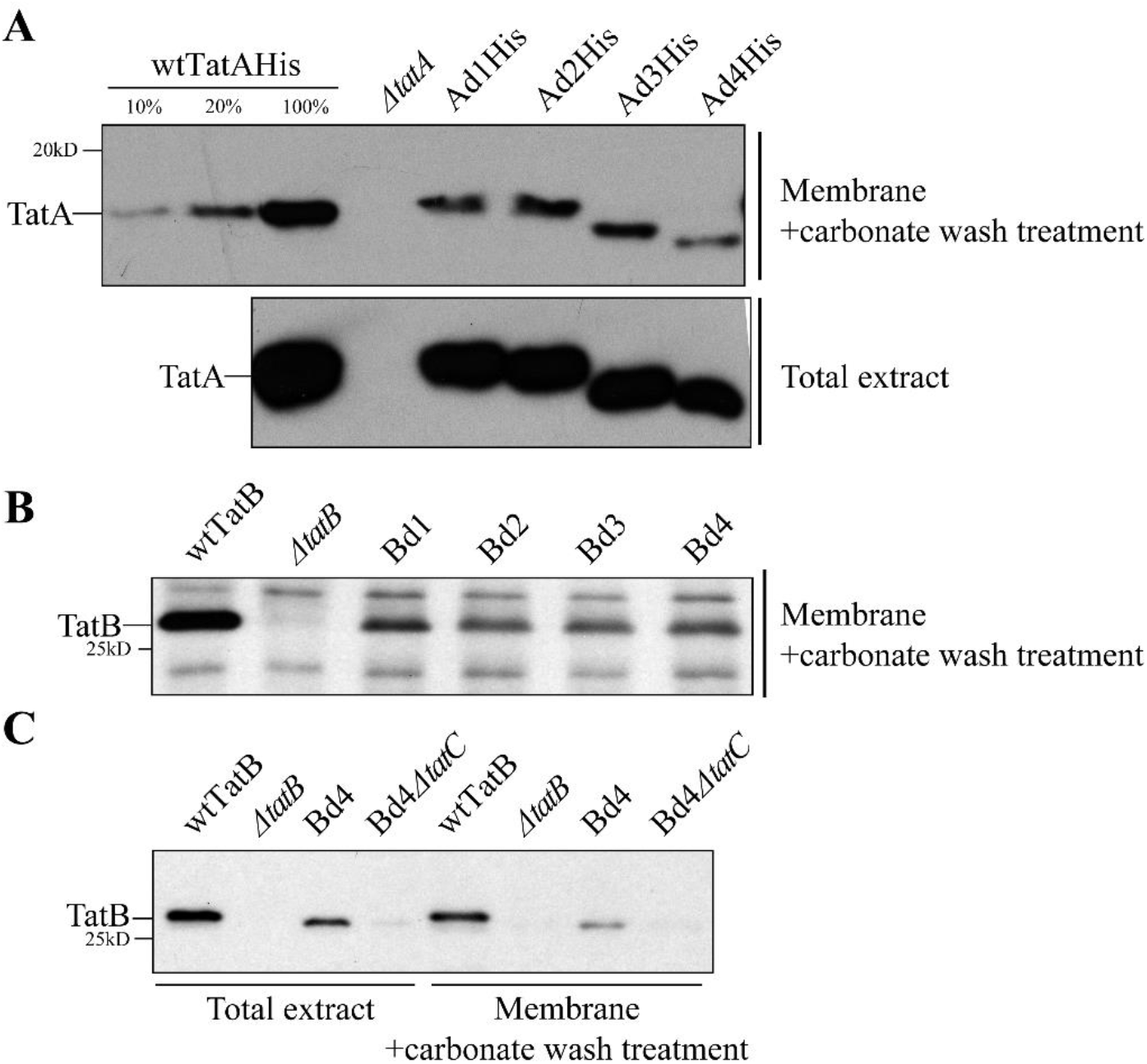
Shortening the TatA and TatB TMH lengths results in lowered membrane stability. **(A)** Assessment of membrane stability in TatA deletion mutants. Total cell extract of C-terminal 6X His-tagged wild-type TatA (wtTatA-His), *ΔtatA*, and TatA deletion mutants (Ad1-4His) were subjected to fractionation. Membrane fractions (upper gel) were recovered from total cell extracts (lower gel) and washed with 100 mM Na_2_CO_3_ to remove the portion of TatA which failed to stably embed in the membrane. Immunoblots using anti-His antibody are shown. Lanes 1-3, serial dilution of membrane-embedded TatA in wild-type cells. (**B**) NaCO3-washed membrane fractions isolated from cell extracts of wtTatB, *ΔtatB*, and TatB deletion mutants (Bd1-4). Immunoblot using anti-TatB antibody is shown. (**C**) Na_2_CO_3_-washed membrane fractions (right) isolated from cell extracts (left) of wtTatB, *ΔtatB*, Bd4, and Bd4 in a TatC knockout strain (Bd4*ΔtatC*). Immunoblot using anti-TatB antibody is shown.

In summary, lengthening the TMHs of TatA and TatB did not significantly affect their membrane stability. Oppositely, even though Ad4 TatA and Bd4 TatB could still insert into the membrane, shortening the TMHs diminished the membrane stability of both TatA and TatB mutants. In addition, TatC, by forming a protein complex with TatB, appears to assist the TatB mutants possessing shorter TMHs to stably embedded in the membrane.

### *E. coli* TatA activity is diminished by lengthening or shortening the TatA TMH

As a first pass at assessing the Tat activity of TatA TMH length mutants we examined their growth profiles in SDS-containing media. Two native *E. coli* Tat substrates, AmiA and AmiC, facilitate cell wall modelling, and a defect in transporting those substrates results in sensitivity of the cell envelope to SDS in the media (Ize et al., 2003). Accordingly, growth in SDS-containing media is a convenient indicator of whether cells have at least a minimal Tat transport capability. Table 1 summarizes the survival ratio after 5 hours growth on SDS-containing LB media for our TMH length mutants; individual survival ratio curves are shown in Supplemental Figure 2. A number of notable points are seen in this table.

First, out of all the members of the ANa group, only ANa1 could grow in LB medium with 5% SDS and the rest lost their ability to grow with as low as 1% SDS (Supplemental Figure 2). In contrast, both A8Qa1-4 and ACa1-4 mutant groups were able to grow in LB medium with 5% SDS, while neither A8Qa5 nor ACa5 could survive in the same media. For the TatA deletion group, surprisingly, all the Ad mutants except the Ad4 could grow in LB medium with 5% SDS. Specifically, the Ad3 mutant, whose TMH only contains 12 amino acids, still retained the ability to grow in LB medium with 5% SDS.

While the SDS growth assay is convenient, it is not obvious how it scales with absolute Tat activity. To examine our mutants more quantitatively, we monitored the transport of a native Tat substrate, SufI, to the periplasm 2.5 hours after induction with IPTG in the presence of TatA mutants in a TatBC background. This assay, while still essentially an end-point assay, has the potential to provide a more fine-grained view of Tat activity than the SDS growth assay. The results of this *in vivo* transport assay, shown in Figure 5 as representative gels from two experiments, display many of the features observed in the SDS growth assay. For instance, ACa1, 2 and 4 mediated SufI transport (Figure 5B) in addition to growth on SDS, as is the case also with the deletion mutants Ad1, Ad2, and Ad3 (Figure 5C). Neither ACa5 nor Ad4 showed SDS growth or SufI *in vivo* transport (Figure 5B and 5C). However, it is clear that none of the mutants that could grow on SDS transported SufI as efficiently as did the wild-type TatA. As an example, see the A8Qa mutants in Figure 5A or any of the mutants in Figure 5B and 5C. Thus, the *in vivo* SufI transport assay leads to the conclusion that either lengthening or shortening the TatA TMH results in loss of Tat transport efficiency, suggesting that there is a functional reason for its 15 amino acid length.

**Figure 5.**
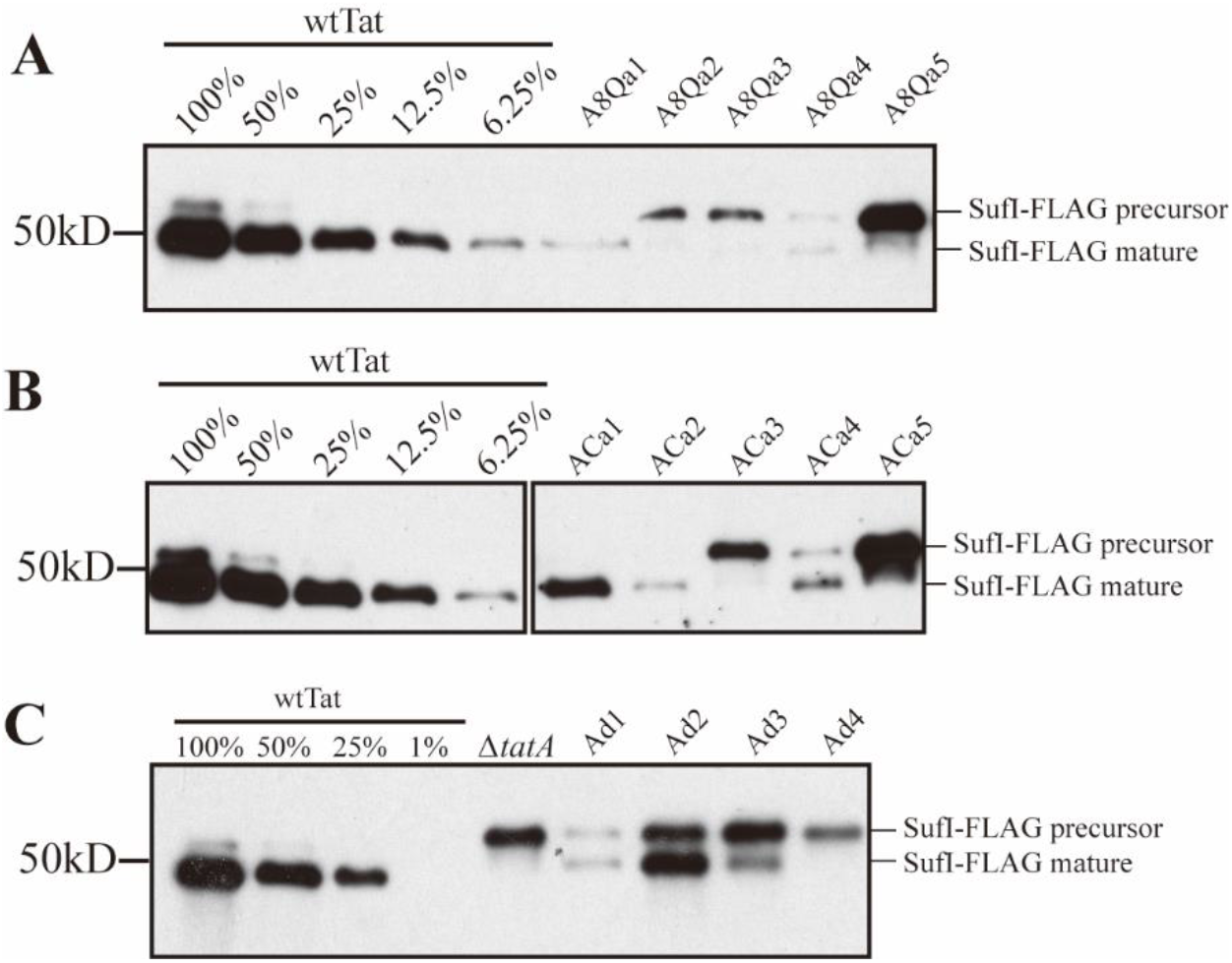
*In vivo* transport of SufI in TatA TMH length mutants. Periplasmic fractions were prepared from wild-type Tat (wtTat), TatA knockout mutant (*ΔtatA*), and TatA mutant cells transporting FLAG-tagged SufI (SufI-FLAG) 2.5 hours after induction with 1mM IPTG. The number of cells used in periplasmic extraction was normalized based on OD600, and immunoblots were developed using anti-FLAG antibody. Precursor and mature forms of SufI-FLAG are labeled. (**A**) transport of SufI-FLAG in the AQ8a group (AQ8a1-5); (**B**) transport of SufI in the TatA C-terminal addition group (ACa1-5); (**C**) transport of SufI-FLAG in the TatA deletion group (Ad1-4).

An even more fine-grained assessment of Tat transport activity is offered in pulse chase assays. Here, kinetics of transport can quantitatively indicate the true extent to which the TMH length mutants operate compared to the wild type. As seen in Figure 6B and 6D, whilst the wild-type TatA transported SufI with initial velocity (V_0_) of 77.1˟ 10^-3^ min^-1^, the next best C-terminal addition mutant, ACa1, operated with a rate constant only 19% of that value. The best performing deletion mutant, Ad2, operated with a rate constant 48% of the wild type (Figure 6C and 6D). Here, when a true quantitative comparison of the TatA TMH length mutants are undertaken, it can again be concluded that the 15-residue wild type length appears to be tuned for optimal activity.

**Figure 6.**
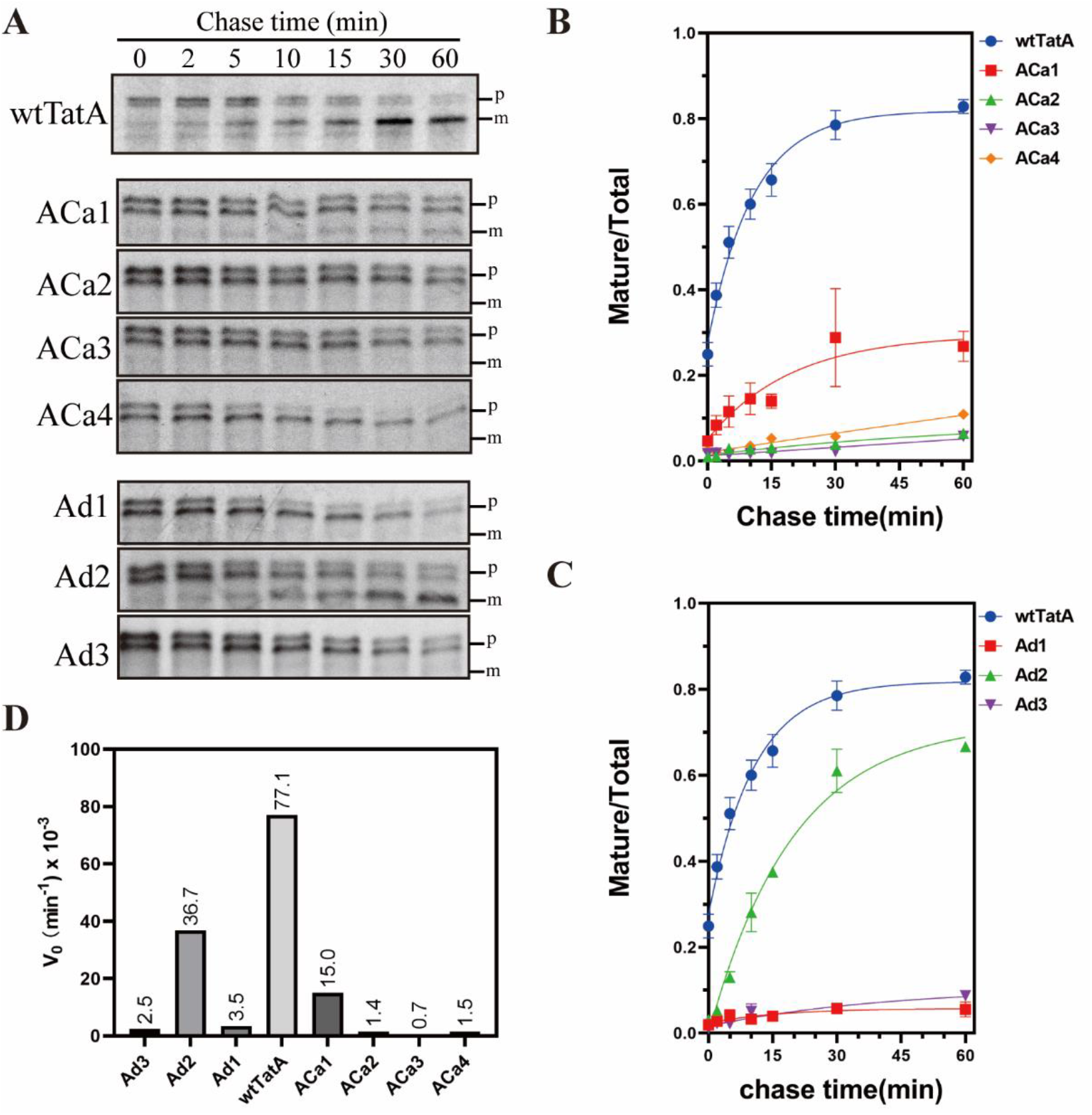
Pulse-chase assays with TatA TMH length mutants. **(A)** SufI transport activity supported by TatA addition and deletion mutants (ACa1-4 and Ad1-3), as monitored by pulse-chase experiments. Autoradiograms developed from 8-16% acrylamide gels are shown; gels are representative of at least 2 biological replicates. p, precursor; m, mature SufI. **(B,C)** Quantitation of the gels in panel A. The ordinate represents the ratio of intensities of the mature bands divided by the intensities of the mature + precursor bands. Data were fitted to a rising exponential model, and the corresponding initial velocity (V_0_) are shown in **(D).**

### Overexpression of Ad3 and Ad4 blocks Tat transport

The majority of TatA is known to be recruited to the TatBC complex after the complex binds to the precursor signal peptide (Alcock et al., 2013; Dabney-Smith et al., 2006, p. 4; Mori and Cline, 2002; Rose et al., 2013). Since the optimal number of TatA is reported to be ∼20-fold higher than the TatB and TatC in the translocon (Celedon and Cline, 2012; Leake et al., 2008), insufficient TatA in the membrane is a possible reason for lowering of the overall transport rate when the TMH is shortened below 15 amino acids.

To determine if the lower abundance of the TatA in the Ad mutants caused the decrease in the Tat activity, wild-type TatA, and the Ad1 – Ad4 TatA mutants were overexpressed on a separate plasmid which was induced by arabinose. Both the wtTatA, and the first three deletion mutants, Ad1, Ad2 and Ad3, and less so for Ad4, accumulated to higher levels after overexpression (Figure 7A). This resulted in transport of a slightly higher amount of SufI-FLAG to the periplasm in the wild type and the first two TatA deletion mutants, Ad1 and Ad2 (Figure 7A). In contrast, SufI-FLAG was not transported when the shortened TatA mutants Ad3 and Ad4 were induced by arabinose, even though they accumulated to levels exceeding or just below that of wtTatA, respectively. When a gradual increase of arabinose concentration was applied to induce increasing amounts of Ad3 TatA, the amount of mature SufI-FLAG was decreased in a dose-dependent manner, indicating that overexpressed Ad3 TatA blocked Tat transport (Figure 7B). Furthermore, when Ad3 and Ad4 TatA mutants were overexpressed in wild-type Tat background, they inhibited SufI-FLAG transport even in the presence of fully functional wtTatA (Figure 7C). These experiments demonstrate that while overexpressing Ad1 and Ad2 TatA could slightly improve the Tat transport activity, overexpressing Ad3 and Ad4 TatA exhibited a dominant negative-like effect. They also show that the relative low abundance of Ad3 and Ad4 TatA in the membrane is not the reason for the low Tat activity seen in Figure 5 and 6.

**Figure 7.**
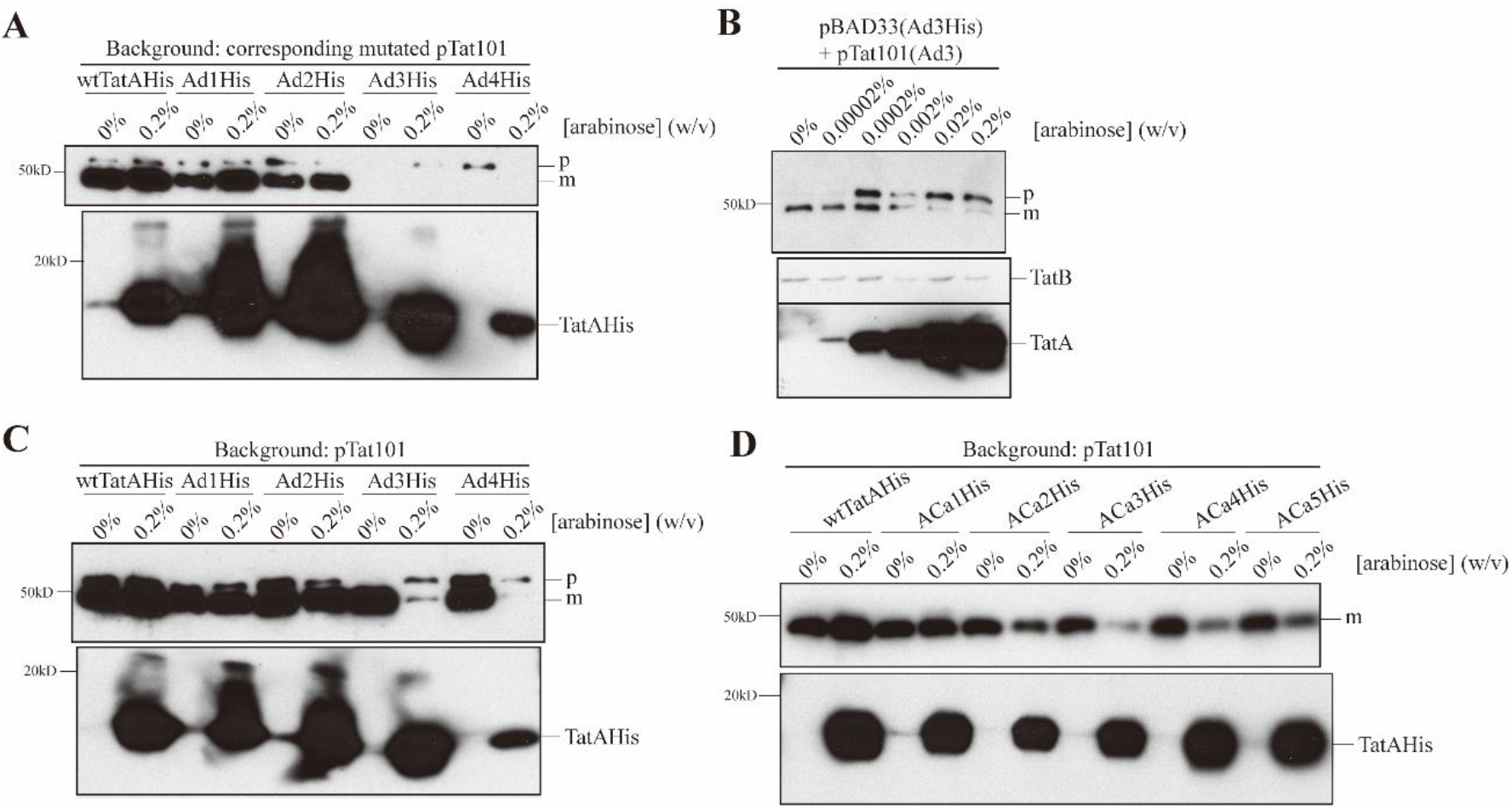
Effect of overexpression of TMH-shortened TatA on SufI transport. (**A**) *In vivo* transport of SufI utilizing indicated mutants. Indicated cells were grown in the presence of indicated [arabinose] for overexpression (from pBAD33) and with the same mutant constitutively expressed as per prior experiments (from pTat101). When the cells reached OD600 of 0.7 preSufI-FLAG was induced by addition of IPTG and then harvested and periplasm prepared therefrom after an additional 2.5 hours. The number of cells used in the periplasm extraction was normalized based on OD600. An immunoblot using anti-FLAG antibody is shown (upper gel). Membrane fractions were isolated carbonate-washed, and samples were then subjected to immunoblotting using the His antibody probing for TatA (lower gel). (**B**) The experiment of panel A was performed with Ad3 only at the indicated arabinose concentrations, and the immunoblot in the lower gel was additionally probes with anti-TatB as a loading control. (**C**) The experiment of panel A was performed with wtTatA expressed as in previous experiments from pTat101 whilst the indicated TatA TMH length mutants were overexpressed from pBAD33 (when arabinose was present). (**D**) *In vivo* transport of SufI with overexpressed ACa group TatA under pTat101 – DADE-A background. Upper gel, periplasm fraction. Lower gel, membrane fraction after sodium carbonate wash. Experimental conditions were essentially same as (**C**). p, precursor; m, mature SufI-FLAG.

A similar dominant negative effect was seen when the ACa mutants were overexpressed in the presence of wild type Tat A (Figure 7D). It can be seen that none of ACa group mutants exhibited high transport activity when overexpressed with 0.2% arabinose. The ACa2-5 mutants especially displayed significantly lower transport when overexpressed with wtTat, even though their effects were not as severe as with the Ad3 and Ad4 mutants (Figure 7C). These experiments show that overexpression of both the Ad and ACa group mutants did not lead to improved Tat transport activity, but rather blocked transport in the more severe cases.

### TatA mutants with shortened TMHs compromise membrane integrity

It has been previously demonstrated that the TatA TMH (without the APH) can compromise membrane integrity in IMVs (Hou et al., 2018). In our study, we asked whether a similar effect would be seen with our TatA mutants which have shortened TMH. Wild-type TatA, Ad TatA or ACa TatA was overexpressed with either constitutively expressed TatBC or in the DADE-Astrain (*Δtat*) background. Relative membrane abundance was also tested by immunoblot (Supplemental Figure 3). Acridine orange was used to detect the ΔpH across the IMVs membrane (Figure 8). Lower ΔpH was interpreted as an increased proton leak.

**Figure 8.**
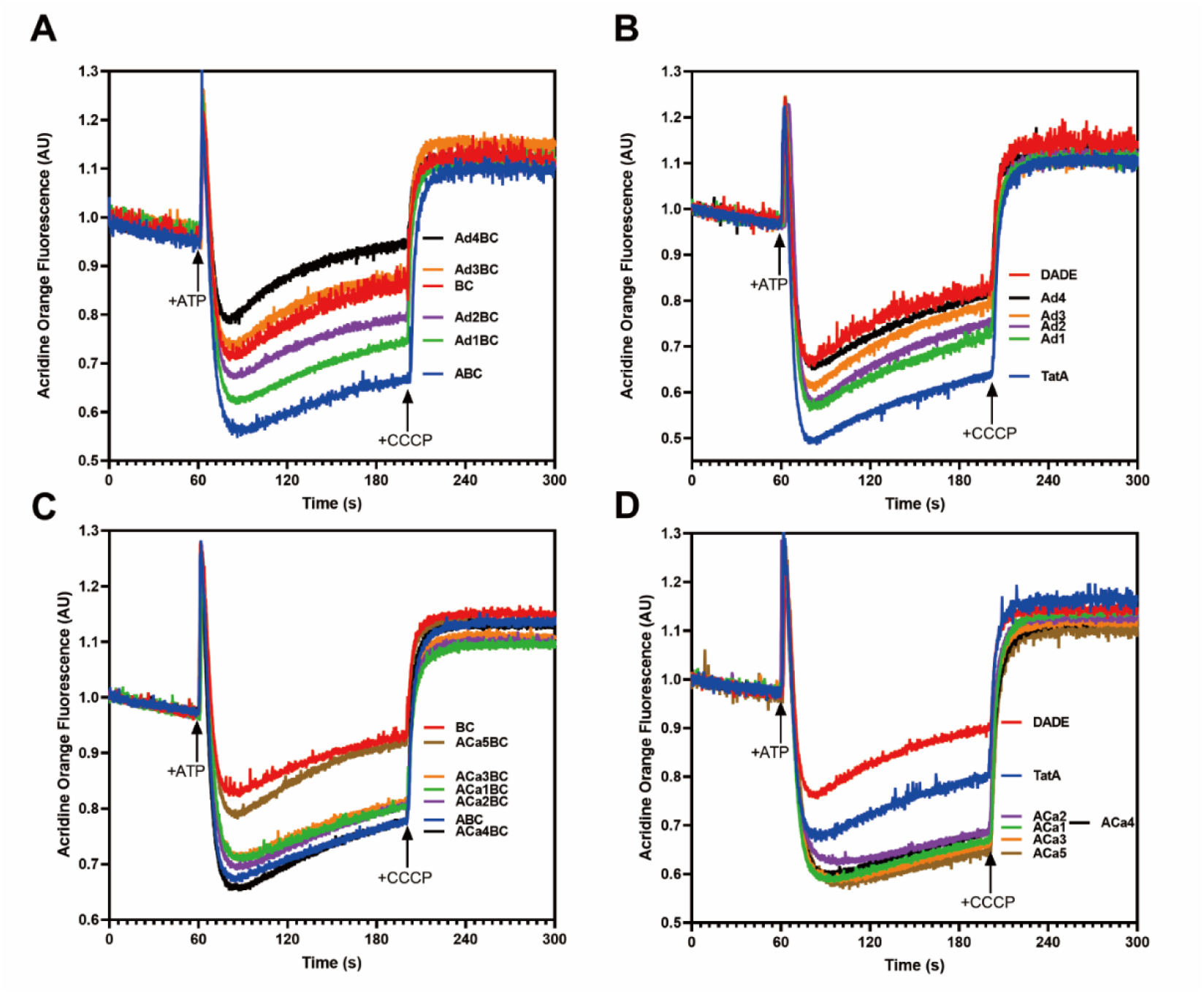
Shortened TatA TMH can cause membrane leakage. The ΔpH developed across IMVs was monitored by quenching of acridine orange. 4 mM ATP was added at 60 sec to generate the ΔpH; 10 μM CCCP was added at 200 sec to dissipate the ΔpH, leading to fluorescence recovery. **(A,C)** ΔpH in TatBC IMVs alone or with the corresponding TatA variants. **(B, D)** ΔpH in DADE (complete Tat knock out) IMVs. AU, arbitrary units.

In these experiments we found first, that wtTatA actually assisted in maintaining membrane integrity, which is counterintuitive since TatA possesses a short TMH. Second, Figure 8A shows that the membranes developed a progressively lower pH gradient (less quenching of the fluorescence signal after adding ATP) when the TMH of TatA was progressively shortened. This result was not related to relative TatA abundance (Supplemental Figure 3). In addition, Ad4 TatA, which showed no Tat activity, caused the highest membrane leakage compared to TatABC or TatBC IMVs. Third, no such effect was observed in TMH lengthened ACa group mutants, with the exception that ACa5BC IMVs displayed similar membrane leakage as TatBC IMVs (Figure 8C). These effects were manifested in the absence (Figure 8B and 8D) of TatBC as well. The Ad4 TatA mutant displayed a similar membrane leakage effect as the no Tat strain. In contrast, the ACa mutants, which possess longer TMHs than wtTatA, displayed much lower membrane leakage than the wtTatA (Figure 8D). We conclude from these experiments that, longer TatA proteins can assist in maintaining membrane integrity in the IMVs, while shorter TatA TMHs resulted in a loss of membrane integrity. The Ad4 TatA in the presence of TatBC caused the highest membrane leakage among all the mutants tested. Such result suggests a general increase in membrane instability when shortening the TMH, which explains why the shorter TMH TatA mutants exhibited lower transport rates than wtTatA in spite of their increased hydrophobic mismatch.

### *E. coli* TatB tolerates a wider range TMH lengths than does TatA

As previously described, the SDS growth assay was conducted with the TatB mutants as well. For TatB addition group, the BNa mutants survived in LB medium with 5% SDS, although BNa5 displayed a lower survival ability compared to other BNa mutants (Table 1). Gradual decrease in Tat activity was nonetheless observed in both *in vivo* transport assays (Figure 9A) and pulse-chase experiment (Figure 10A, B and D). The BNa1, BNa2 and BNa3 mutants retained approximately 58%, 13% and 3.5% of the wild-type TatB V_0_, respectively. In contrast, none of B8Ea and BCa group mutants exhibited SDS tolerance, indicating that no Tat transport occurred in those mutants.

**Figure 9.**
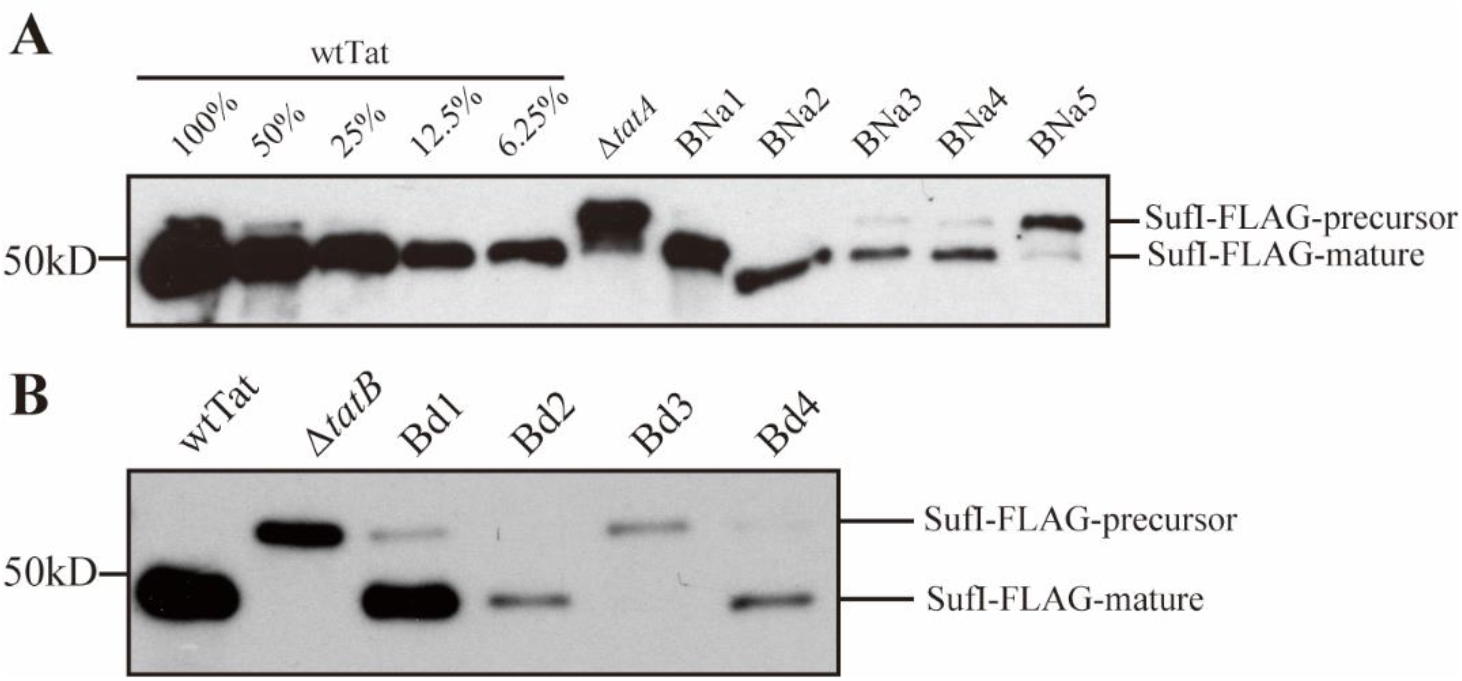
*In vivo* transport assays with TatB mutants. Experiments were performed essentially as in Figure 5 but substituting the TatB TMH length mutants that grew on SDS for the TatA mutants of that figure. (**A**) Transport of SufI-FLAG in the BNa TatB addition group. (**B**) Transport of SufI-FLAG in the Bd TatB deletion group.

In contrast to the TatB addition mutants, a special pattern in the TatB deletion group mutants in terms of SDS tolerance was identified. The Bd1, Bd2 and Bd4 mutants retained sufficient Tat activity to grow in LB medium with 5% SDS, while the Bd3 mutant did not. However, pulse chase experiments with the Bd group indicated that only approximately 5% transport activity was retained in the Bd1, Bd2 and Bd4 mutants (Figure 10C and D). This speaks to the coarseness of the correlation between growth on SDS-containing media and the absolute Tat activity (see Discussion).

**Figure 10.**
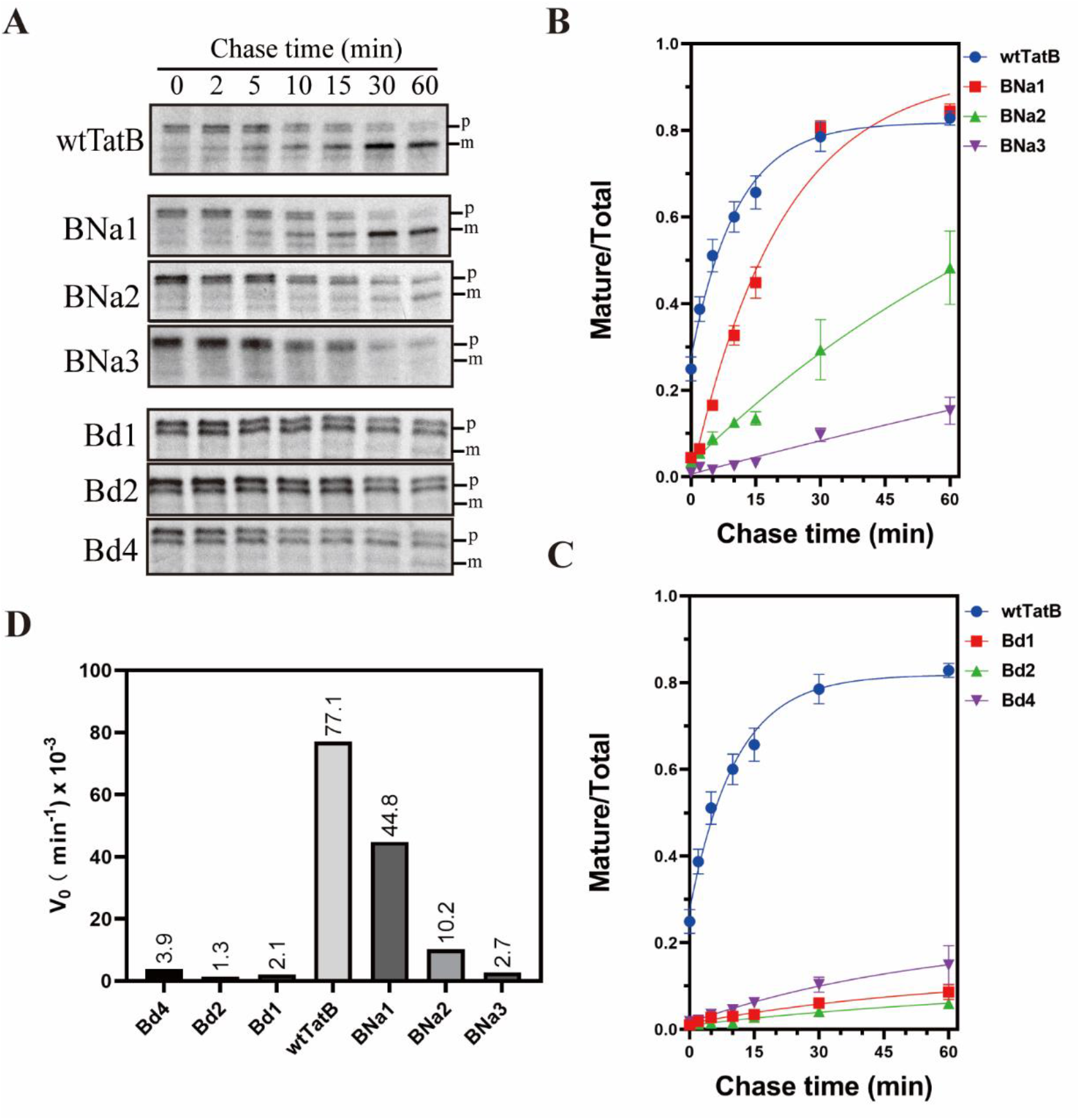
Pulse-chase assays with TatB TMH length mutants. Experiments were performed essentially as in Figure 6 but substituting the TatB TMH length mutants that grew on SDS for the TatA mutants of that figure. (**A, B, C, D**) SufI transport activity supported by TatB addition and deletion mutants (BNa1-3 and Bd1, 2, 4), as monitored by pulse-chase experiments. Details as described in the Figure 6 legend.

To summarize, these results indicate an acceptable range from eleven to twenty amino acids in the TatB TMH, which is wider than the range proposed from the TatA mutants. As with TatA, a change in the length of the conserved hydrophobic region in the TatB TMH is not favored. Even though an overall longer TMH inhibits the TatB function, if the length of the hydrophobic region is preserved (i.e., adding residue from the N-terminus), its negative impact is not as severe when the length of the hydrophobic region is altered.

## Discussion

In this study, the effect of the hydrophobic mismatches in the Tat translocon was examined by modifying the length of the TMHs in *E. coli* TatA and TatB. Growth on SDS-containing media, and *in vivo* transport and pulse-chase assays were conducted to evaluate the Tat transport activity in the TMH length mutants comprehensively. The results showed that while both TatA and TatB can tolerate some length modification in their respective TMHs, none of the modified mutants transported Tat substrates as well as the wild-type strain. Interestingly, TatA and TatB exhibited different acceptable lengths of their TMHs, with TatA tolerating a TMH 11-19 residues long and TatB tolerating a length between 10 and 20 residues. Further comparison of the transport rates between the addition and deletion mutants revealed different behaviors of the TatA and TatB TMH length mutants, perhaps reflecting their different roles in the Tat transport mechanism.

It is important to stress that TMHs both lengthened or shortened did not support protein transport as well as the wild type 15 amino length TMHs of both TatA and TatB. This, along with the observations that the 15-residue length of these TMHs is conserved (Figure 2 A, B), and moreover, this length is relatively rare in single-pass membrane proteins (Figure 3B), suggests that this particular hydrophobic mismatch is evolutionarily tuned for maximum activity. We are not the first to propose that this plays a role in the mechanism of Tat protein transport (Hou et al., 2018; Rodriguez et al., 2013).

One complicating issue in interpreting the fall off in activity of the TMH length mutants is the fact that amino acids in these TMHs interact with other residues, both in the same polypeptide and in TatC. Co-evolution analysis predicted as many as ten interactions between the TatA TMH residues and those on helices 5 and 6 in TatC (Alcock et al., 2016). Were we to add or delete additional amino acids directly into the middle of the TatA TMH, for instance, then we could expect to disrupt such interactions as positions of amino acids arrayed on the interacting face of the helix would have been altered. To minimize this potential complication, we added residues to the extreme N- and C-termini of the TMHs. Addition or deletion of residues form the C-terminus would be expected to keep the presentation face of the TMH intact, while possibly changing its orientation with respect to the APH. This is illustrated in Figure 11, which shows that in the wild type proteins the APHs of both TatA and TatB are displace approximately 100 – 120 degrees from their respective polar amino acids. The importance of this relative displacement is not obvious, as growth on SDS-containing media cannot be predicted by proximity in the helical wheel projection of the APH to the polar amino acid. Nonetheless, amino acid additions at the extreme C-terminus would be expected to maintain the presenting face of the TMH, and in the case of the ACa mutants, all interacting residues should be intact and in position.

**Figure 11.**
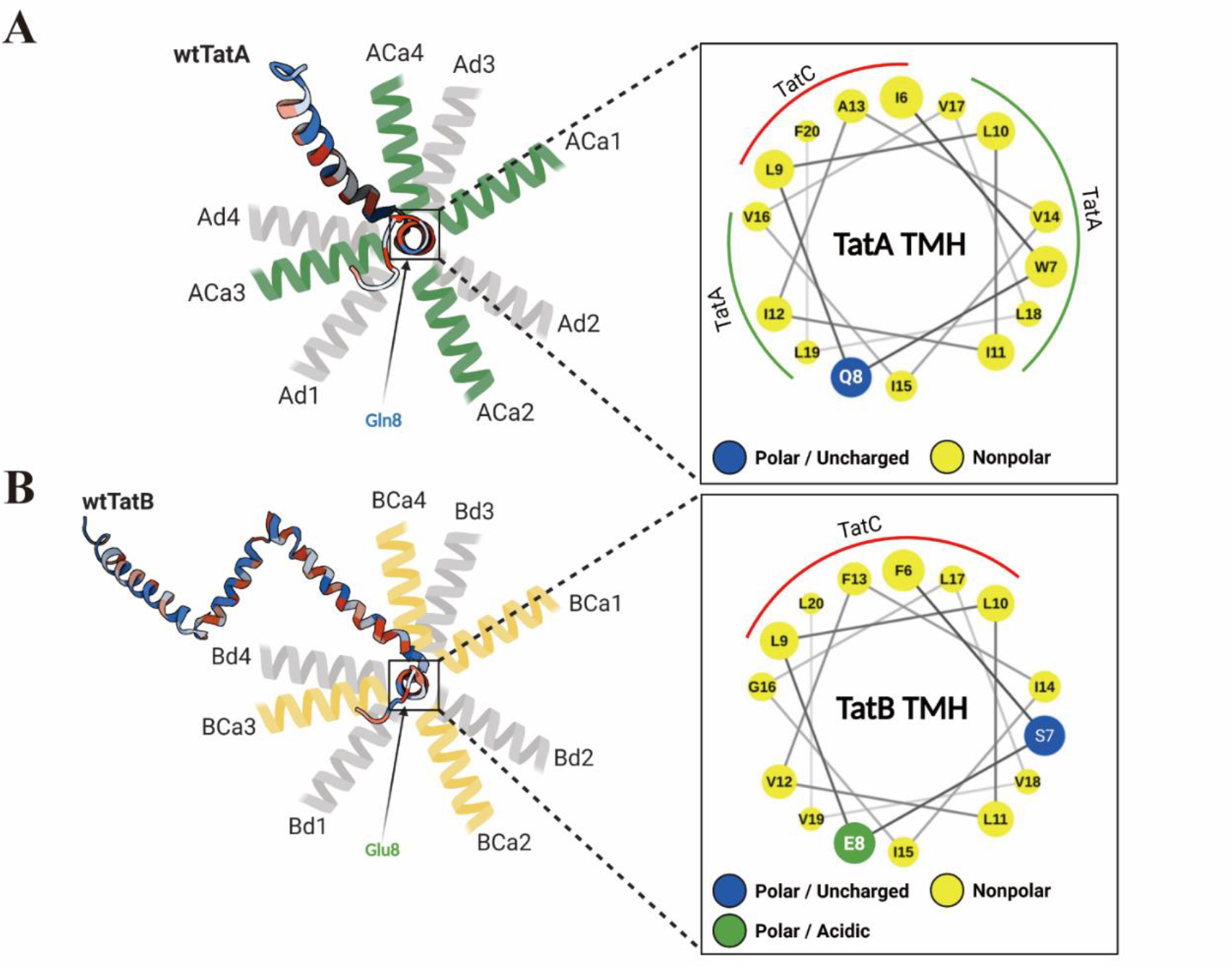
Possible APH orientations change in TatA and TatB TMH mutants. Predicted TatA (A) or TatB (B) APH orientations are illustrated from the top view. The projected structures of wtTatA (ID: 2MN7) and wtTatB (ID: 2MI2) showing the possible orientation of the APHs with respect to non-variant polar amino acid positions are shown on the left. On the right is shown a helical wheel projection (Mól et al., 2018) showing the potential Tat subunit interaction faces.

A less complex situation is expected when amino acids are added to the N-terminus of the TMH. In this instance, only the extreme N-terminus of the protein should rotate relative to the TMH, with the only potential contact disrupted being the 5^th^ amino acid in either TatA (G5) or TatB (S5); all other residues are intact and in position. Still, one possible effect of such N-terminal addition mutants could be that they push the polar amino acid at position 8, which was shown to be functionally important (Greene et al., 2007), deeper into the middle of the membrane.

We further considered the possibility that extended-TMH TatA variants do not support protein translocation because they are too long to fit comfortably into the TatBC complex. However, this seems an unlikely explanation because the ACa mutants exhibit a dominant negative effect when overexpressed in the presence of wtTatA. In that experiment, we would not expect the lengthened TatA mutant to displace the wtTatA if it does not fit into the TatA binding site in the first place. Second, it has been reported that TatA and TatB might switch contact positions with TatC in the presence of substrates (Johann et al., 2017). In this study we show that TatB can tolerate a larger range of TMH lengths than TatA, especially for the BNa mutants (Figure 10). If the receptor complex could only fit a TatA or TatB TMH of a particular size, we might expect identical inhibitory behaviors of the TMH length mutants of these two proteins; we did not. Accordingly, we look to other explanations for their inhibition of protein transport beyond an inability to bind to the TatBC complex.

It is perhaps unexpected that the deletion mutants as short as 11 residues were still able to take up residence in the membranes and resist extraction by carbonate. However, examples of short peptides integrating into both artificial and biological membranes are known from the literature (Ulmschneider et al. 2018; de Planque et al. 2003; Jaud et al. 2009) Additionally, several studies have shown that soluble TatA is able to be functionally incorporated into thylakoid membranes (Aldridge et al., 2012; Dabney-Smith et al., 2006, 2003; Frielingsdorf et al., 2008; Hauer et al., 2013; Pettersson et al., 2021). The TatB Bd4 mutant loses its membrane location in the absence of TatC (Figure 4C), which points to a stabilizing interaction between the short TatB and TatC. However, the TatA Ad4 mutant appears to embed in the membrane on its own. This is demonstrated not only by their retention in the membranes after carbonate washing, but also by the dominant negative-like effect on SufI transport of Ad3 and Ad4 overexpression in the presence of wild type TatA, and by their ability to cause proton leakage in IMVs.

Our analyses of our TMH length mutants revealed an important technical point about assays for Tat activity. Given that the ANa mutants (except ANa1 mutant) were unable to grow in the presence of SDS, we were surprised that the ACa mutants could indeed grow on the same SDS-containing media. It was only when we performed kinetic assays that it became clear that the Tat activity in the ACa mutants was quite low. Even the best performing TatA C-terminal addition mutant, ACa1, transported SufI at only about 19% of that of the strain harboring wild type TatA (Figure 6B). Accordingly, the SDS-growth assay, which is an end point assay, does not scale linearly with Tat activity, and generally indicates only the complete loss of Tat function. Even ACa2, which transports SufI at 1.8% the rate of wild type showed wild type-like survivability on up to 10% SDS. Assays of Tat function by monitoring the presence of SufI in the periplasm are somewhat more sensitive to absolute Tat activity, but again, this is an end point assay that may underestimate the negative impact of an altered Tat machinery.

We were surprised to see the different effects of amino acid addition after the polar residues in TatA and TatB. While the amount of SufI found in the periplasm in the TatA A8Qa mutants was generally quite low (Figure 5A), they retained enough activity for growth in the presence of SDS. This contrasted the similar additions in TatB after the E8 residue, which apparently had no remaining Tat activity. This is just one example of many in which lengthening the TMH of TatA produced a different effect than a similar change in the length of the TMH of TatB. It is tempting to attribute this to a proposed fundamentally different function of these two proteins in the mechanism of Tat transport.

We also found that lengthening additions were sensitive to their location relative to the polar amino acids in these TMHs. In both TatA and TatB additions made at the extreme N-terminus of the TMH had different effects than those made just two residues further into the helix, after the polar amino acids. Additions made after the 7^th^ amino acid, but just before the 8^th^ polar residue had the same growth profile in SDS as the N-terminal additions (Supplemental Figure 2). The reason that the addition phenotypes switch around the polar residues remains to be elucidated, but possibilities include disruption of the helix interacting faces, and changes of the depth of the polar amino acids in the membrane.

How do these TMH length mutants inform us about the mechanism of protein transport on the Tat pathway? We knew already that the 15 amino acid length of the TatA and TatB TMHs are evolutionarily optimized by the fact that this length is so conserved. Our experiments now point to why this is the case. It is clear that changing the TMH lengths of either TatA or TatB from 15 residues in either direction results in a decline in overall Tat activity. One could argue that this decline in the TatB TMH mutants might be related to a loss of its ability to interact with TatC correctly, although the advantage conferred by making additions at the N-terminus of the TMH are still in effect. Still, since TatB and TatC form a stable complex with a 1:1 stoichiometry, it is difficult to attribute any function to TatB alone. TatA, on the other hand, has been studied on its own and has been shown to have interesting properties by itself (Alcock et al., 2013; Celler et al., 2013; Gohlke et al., 2005; Hauer et al., 2017; Hou et al., 2018; Zhang et al., 2014a). We interpret the detrimental effects of TatA TMH shortening and lengthening differently. We note that while the Ad2 deletion mutant has considerably better activity than any addition mutants (Figures 5C and 6B), which suggests that the increased hydrophobic mismatch confers some advantage, it still does not perform as well as the wild type. This decline in Tat activity when the TatA TMH is shortened could be attributed in part or in full to a loss of PMF driving force due to induced membrane leakage to protons.

The TatA TMH addition mutants also display poor Tat activity, although we believe there is a different mechanism at work here. We favor the hypothesis that Tat protein transport occurs through transient toroidal pores that form in the membrane as a result of bilayer breakdown in response to localized membrane thinning (Asher and Theg, 2021; Berks, 2015; Brüser and Sanders, 2003; Chen et al., 2003). In this model, the hydrophobic mismatch set up by the Tat subunits confers a natural advantage toward membrane thinning, and this would be intensified by the binding of more TatAs with their short TMHs to the active machinery as the latter assembles on demand (Rollauer et al., 2012). Further, extrapolating the effect of the PMF on thinning of the thylakoid membrane (Johnson et al., 2011, 2011; Kirchhoff et al., 2011; Murakami and Packer, 1970) to the *E. coli* plasma membrane as well provides an immediate mechanism for coupling the PMF to Tat protein transport. Alterations in membrane thickness as a critical determinant of membrane protein function and/or assembly has recently been noted by others (Chen et al., 2017; He et al., 2020; Iadanza et al., 2020; Kreutzberger et al., 2019; Pleiner et al., 2020; Wu et al., 2020). In this context, one might expect that hydrophobic mismatch in the Tat translocation machinery would play a key role in Tat transport. Accordingly, the tuning of the TatA and TatB TMHs to 15 amino acids might be seen as an evolutionary compromise of selecting a hydrophobic mismatch that is not so severe as to cause detrimental ion leakage, but still sufficient to allow the membrane to thin enough under physiological conditions to the point of toroidal pore formation.

## Materials and Methods

### Strain and plasmid construction

*E. coli* strain DADE-A (MC4100, *ΔtatABC, ΔtatE*, arabinose resistance) was used in both *in vivo* and *in vitro* experiments in this study (Wexler et al., 2000). For TatA and TatB variants in pTat101 (pTH19Kr derivative, a low copy plasmid constitutively expressing TatABC) (Kneuper et al., 2012), the indicated deletion and addition mutations were introduced by QuickChange site-directed mutagenesis (NEB, Q5 Site-Directed Mutagenesis Kit). For TatA variants in pBAD33 (a vector with arabinose-inducible araBAD operon) (Guzman et al., 1995), the appropriate *tatA* mutant alleles were amplified from the corresponding constructs in pTat101, which were then assembled into pBAD33 using the Gibson assembly approach (Gibson et al., 2009). The 6XHis tag was inserted into the indicated TatA constructs in pTat101 using the primers TatAhis_F (5’-CACCACCACTAACACGTGTTTGATATCG-3’) and TatAhis_R (5’-ATGATGATGCACCTGCTCTTTATCGTG-3’). For TatA constructs in pBAD33, the *His6* tag was added using the primers pBAD33TatAhis_F (5’-TCACCACCACTAATGGCTGTTTTGGCGG-3’) and pBAD33TatAhis_R (5’-TGATGATGACCCACCTGCTCTTTATCGTG-3’). For *in vivo* transport experiments, construct of pQE80l (SufI-FLAG) was produced as described (Huang and Palmer, 2017). For pulse-chase experiments, constructs pNR14 and pNR42 were as described previously (Sargent et al., 1999; Stanley et al., 2000a). All constructs were confirmed by Sanger sequencing. More detailed information about the plasmids used in this study can be found in Supplemental Table 1.

### Sequence Alignments and Sequence Logo Plots

122 TatA sequences and 60 TatB sequences were downloaded from GenBank (NCBI). Multiple sequence alignment for TatA and TatB, respectively, was performed using MUSCLE (Edgar, 2004). Sequence logos were subsequently generated using RStudio (Ver. 1.3.1073) with the ggseqlogo package (Wagih, 2017).

### Liquid SDS Growth Assay

Overnight cultures grown at 37°C were normalized to an OD600 of 0.1 before adding to the Luria-Bertani (LB) medium containing 0%, 1%, 2%, 5%, or 10% SDS, respectively, to a final optical density of 0.002. After 5 hours of shaking at 37°C, the optical density of the cell suspension at 600 nm was measured. Survival rates of the cells in the LB with corresponding SDS concentrations were calculated by taking the ratio of the optical density of cells grown in LB with indicated SDS concentration to cells grown in the LB without SDS.

### *In vivo* Transport Assay

pTat101 variants were co-transformed with pQE80l(SufI-FLAG) into the DADE-A strain. Cells were first grown overnight at 37°C and were then diluted to an OD600 of 0.06 and cultured in 5 mL of fresh LB medium at 37°C with shaking for 3 hours. SufI-FLAG was induced by the addition of 1 mM IPTG (isopropyl β-D-1-thiogalactopyranoside). Following 2.5 hours of growth at 37°C, cells were harvested and subjected to fractionation. For TatA deletion mutants in pBAD33 variants, plasmids containing indicated mutated *tatA* alleles in pBAD33 were co-transformed with pQE80l (SufI-FLAG), and pTat101 or the corresponding pTat101 variants. Overnight cultures were diluted and sub-cultured in 5 mL of fresh LB medium containing arabinose concentration ranging from 0% to 0.2%. At OD600 ∼ 0.6, cells were induced with 1 mM IPTG. Following 2.5 hours of growth at 37°C, cells were harvested and subjected to fractionation.

### Cell Fractionation

After *in vivo* transport, cells were harvested. The volume of cells in each mutant was normalized based on cell densities such that each sample contained 3 mL of cells with OD600 = 1.5. Cells were centrifuged at 16,000 x *g* at room temperature. Cells were then cooled on ice and fractionated using the EDTA/lysozyme/cold osmotic shock method (Petiti et al., 2017) by applying 80 μL of 1X TES buffer (200 mM Tris-HCl, pH 8.0, 0.5mM EDTA (ethylenediaminetetraacetic acid), 0.5 M sucrose), 3.2 μL of 10 mg/mL freshly prepared lysozyme solution, and 288 μL of 0.5X TES buffer (1X TES buffer diluted twice in water (v/v)), in order. Samples were then incubated at 4°C for 30 min before centrifugation at 5,000 x *g* for 5 min at 4°C. Supernatants were kept as the periplasmic fractions by adding equal volume of 2X SDS sample buffer and were subjected to SDS-PAGE/Western-Blot analyses. For membrane extraction, pellets were then resuspended in 0.5X TES buffer containing 2 mM PMSF (phenylmethylsulphonyl fluoride), 2 mM MgCl2, and 10μg/mL DNase I, followed by 4 cycles of freezing and thawing in liquid nitrogen and centrifugation at 2,000 x *g* at 4°C. For carbonate washed samples, cells were washed in 10 mM Na_2_CO_3_ for 1 hour followed by ultracentrifugation at 120,000 x *g* for 45 min at 4°C. Pellets were kept as the membrane fraction, which were then subjected to SDS-PAGE/Western-Blot analyses.

### SDS-PAGE and Western-Blot

Proteins were separated by SDS-PAGE followed by Western-Blot using anti-TatA, anti-TatB, anti-His tag (Genscript) or anti-FLAG (Genscript) antibodies, depending on the protein samples. *His6,* TatA and TatB were detected by HRP (horseradish peroxide)-conjugated anti-rabbit antibody and SufI-FLAG was detected using HRP-conjugated anti-mouse antibody. Proteins were then visualized using the ProSignal Pico ECL Western Blotting detection kit (Genesee Scientific).

### Pulse-Chase Experiment and Autoradiography

Experimental procedures were derived from (Stanley et al., 2000b). Overnight cultures carrying Tat variants in pTat101, pNR42 and pNR14 were grown in LB media at 30°C. 100 μL of the overnight culture was then added in 3 mL of fresh LB media for sub-culture at 30°C. After 1.5 hours, cells were harvested and normalized such that each sample contained an equivalent of 0.5 mL cells with OD600 = 0.2. Cells were then washed with 1X M9 medium (M9 salt, 0.1 mM CaCl2, 0.002% thiamine, 2 mM MgSO4, and a 0.01% 18-amino acid mix free of methionine and cysteine) to remove excess LB medium and resuspended in 2.5 mL M9 medium. Cells were grown for another hour at 30°C. Subsequently, cells were grown at 42°C for 15 min to induce transcription of T7 polymerase from pNR42. 400 μg/mL rifampicin were added to inhibit the *E. coli* endogenous RNA polymerase, followed by another 10 min of growth at 42°C. Cells were incubated for another 20 min at 30°C. Subsequently, cells were transferred to 37°C until the completion of the experiment. 0.025 mCi of [^35^S] methionine (PerkinElmer Inc. NEG772002MC) was added to 2.5 mL of culture to initiate the pulse process. After 5 min of pulse, cells were chased by adding 750 μg/mL unlabeled cold methionine. A 300 μL sample was taken at each time point, followed by immediate freezing in liquid nitrogen. Samples were then thawed on ice, centrifuged, and resuspended with 50 μl 2X SDS sample buffer, and then subjected to SDS-PAGE and autoradiography. Quantification of the protein bands were carried out using ImageJ software (Schneider et al., 2012).

### Statistical Analysis

Data from the pulse-chase experiments were subjected to statistical analysis. An exponential plateau model, Y = Ym-(Ym-Y0) *exp(-k*x), which is derived from the first-order reaction model, was used to fit the data using GraphPad Prism version 8.2.1 for Windows (GraphPad Software, San Diego, California USA) The ordinate value corresponds to the mature-to-total ratio for each sample, and the abscissa represents time in the pulse-chase experiment. Y_m_ was defined as the maximum value of the mature-to-total (i.e., mature to the sum of the mature and precursor), the initial velocity (V_0_)was obtained from the model (V_0_=Y_m_ * k), with the units of the reciprocal minutes.

### Statistical Analysis of Single-Pass Membrane Proteins

A total number of 9232 single-pass membrane protein sequences from four categories (495 from *E. coli*, 5468 from proteobacteria, 2146 from chloroplasts, and 1397 from mitochondria) were downloaded from Swiss-Prot database (The UniProt Consortium, 2021) by selecting single-pass proteins, followed by selecting the corresponding categories. The transmembrane domains of the proteins were then identified by using the TMHMM Server, v.2.0 (Möller et al., 2001). The output of the expected number of amino acids in the transmembrane helix (TMH) per protein was subsequently collected. The amino acid numbers predicted in the TMHs were rounded to the closest integer, and their relative frequency (i.e., ratio of occurrence to the total number of proteins in the indicated category) was plotted against the TMH length for each category using GraphPad Prism version 8.2.1.

### Proton leakage measurement

Plasmids carrying wild-type or indicated TatA variants with the *His6* tag in pBAD33 were co-transformed with or without TatA in pTat101 separately into DADE-A. IMV preparation was performed as described (Bageshwar and Musser, 2007). Acridine orange fluorescence-quenching assays were performed on a Fluorolog-3 spectrofluorometer (HORIBA Scientific, model No. FL3-22). 50 μl IMVs (final A280=0.375) were added to a 2 ml reaction master mix containing 1X TE buffer (25 mM MOPS, 25 mM MES, 5 mM MgCl2, 50 mM KCl, 200 mM sucrose and 57 μg/ml BSA, pH = 7.0), 2.9 mM phosphocreatine, 0.29 mg/ml creatine kinase, and 2 μM acridine orange. The reaction mix was kept on ice before the measurement. Before the measurement, the mixture was first incubated in a 3 ml cuvette at 37°C for 5 min with slow stirring for temperature equilibration. Acridine orange fluorescence was recorded at λex = 494 nm (slit = 1 nm) and λem= 540 nm (slit = 5 nm) every 1/10 second. 20 μl of 400 mM ATP (4 mM final concentration) was added into the cuvette at 60 seconds, and 4 μl of 5 mM CCCP (10 μM final concentration) was added to the cuvette to dissipate the proton gradient at 200 seconds.

## Acknowledgements

We would like to thank Tracy Palmer and Jon Cherry for their generous gifts of materials, as well as for their comments and discussions on this project. We also thank Ben Berks and Thomas Br<u>ü</u>ser for their many insightful comments on this work. Thank you to Iniyan Ganesan for providing the illustration in Figure 1. We gratefully acknowledge support from the Division of Chemical Sciences, Geosciences, and Biosciences, Office of Basic Energy Sciences of the U.S. Department of Energy through Grant DE-SC0020304 to SMT.

## Competing interests

The authors have no competing interests to declare.

## Supplemental Information

**Supplemental Figure 1.**
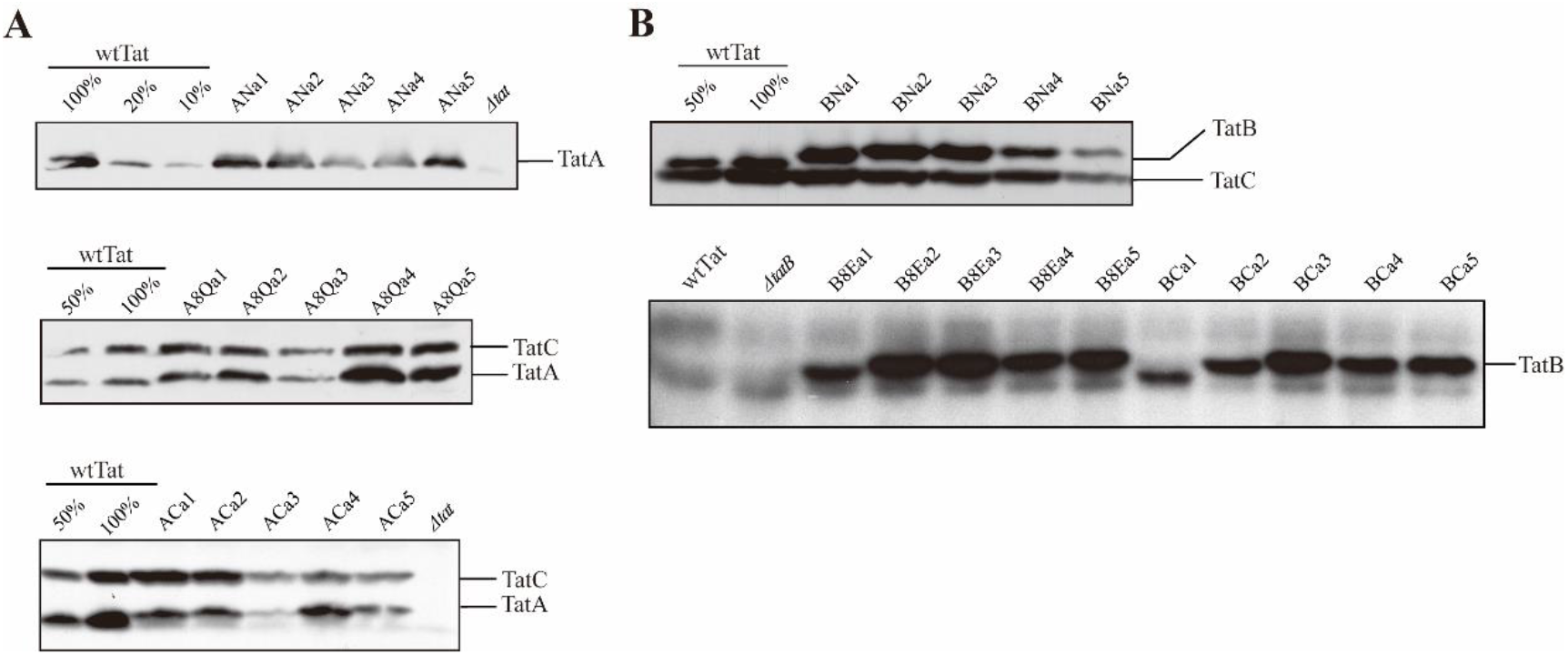
Membrane stability of TatA and TatB in the corresponding addition mutants. (**A**) Assessment of membrane stability in TatA addition mutants. Shown are immunoblots of carbonate-treated membrane fractions from extracts of wild-type Tat (wtTat), *ΔtatA*, and TatA addition mutants lengthened at three different locations (ANa1-5, AQa1-5, and ACa1-5) developed with anti-TatA antibody, without (upper) or with anti-TatC antibody (lower). (**B**) Shown are immunoblots of carbonate-treated membrane fractions from extracts of wild-type Tat (wtTat), *ΔtatB*, and TatB addition mutants lengthened at three locations (BNa1-5, BEa105, and BCa1-5) developed with anti-TatB antibody with (upper) and without (lower) anti-TatC antibody.

**Supplemental Figure 2.**
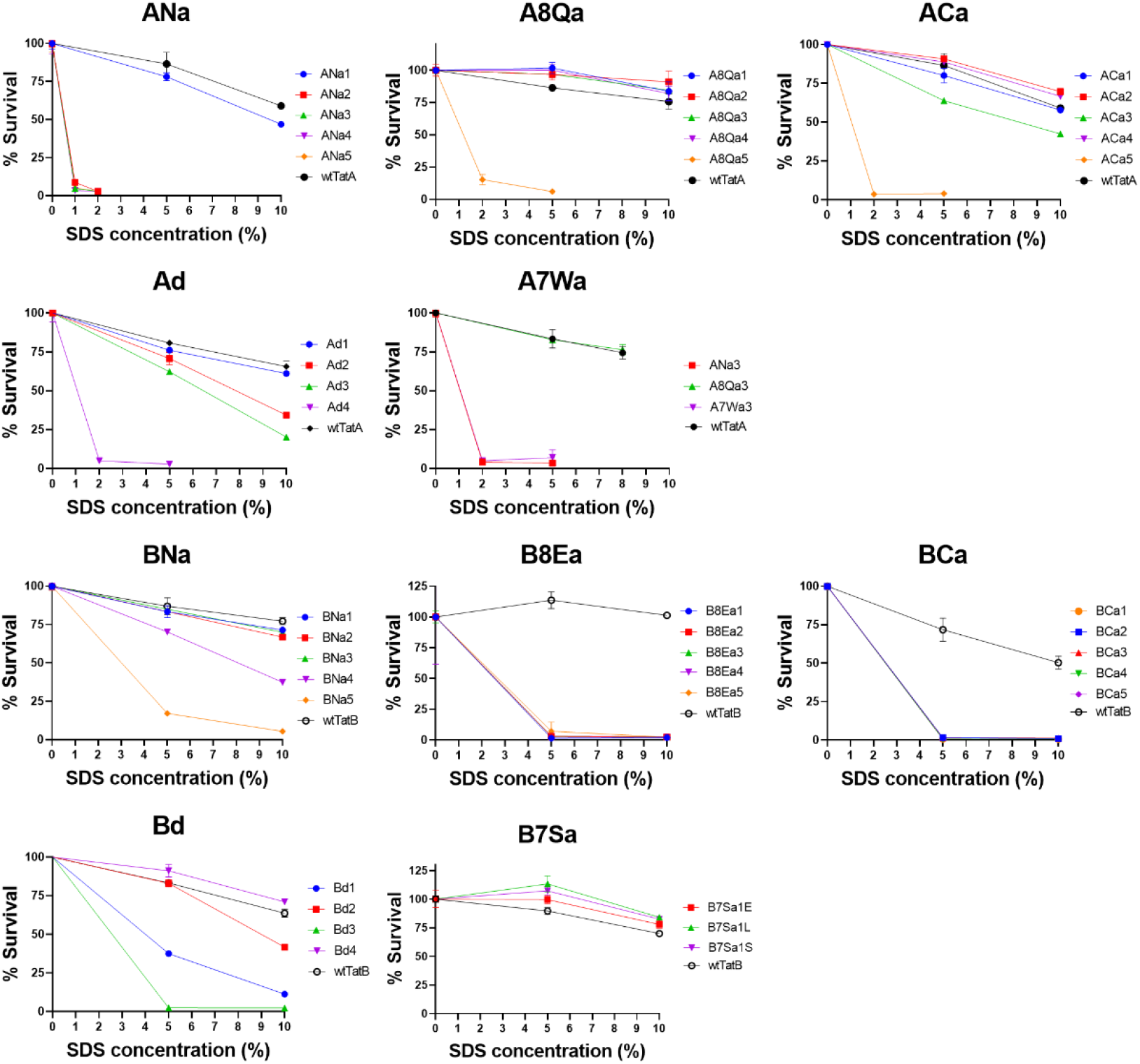
Growth performance of the TatA and TatB TMH length mutants in the presence of SDS. wtTatA and wtTatB act as the positive controls. Survival ratios, displayed in percentages, were obtained by computing the ratio of the optical density of the cells grown for five hours in the indicated SDS concentration to the cells grown without SDS.

**Supplemental Figure 3.**
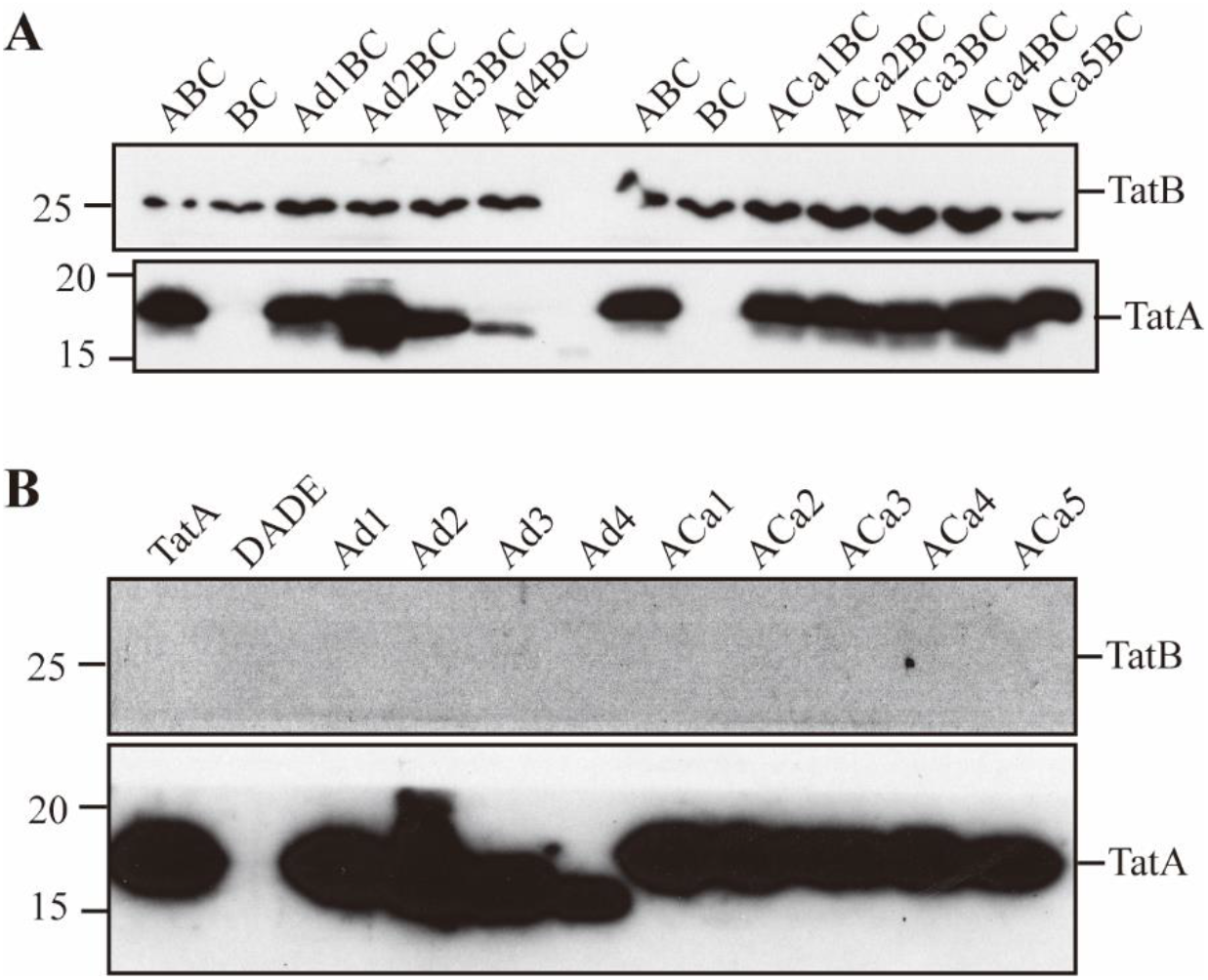
Confirmation of the expression of mutated TatA in TatBC IMVs and DADE IMVs. (A) TatBC IMVs or (B) DADE IMVs alone or with the corresponding His-tagged TatA variants were subjected to immunoblotting using anti-His antibody probing for TatA. TatB, which was detected by TatB antibody, served as a loading control for this experiment.

**Supplemental Table 1.**
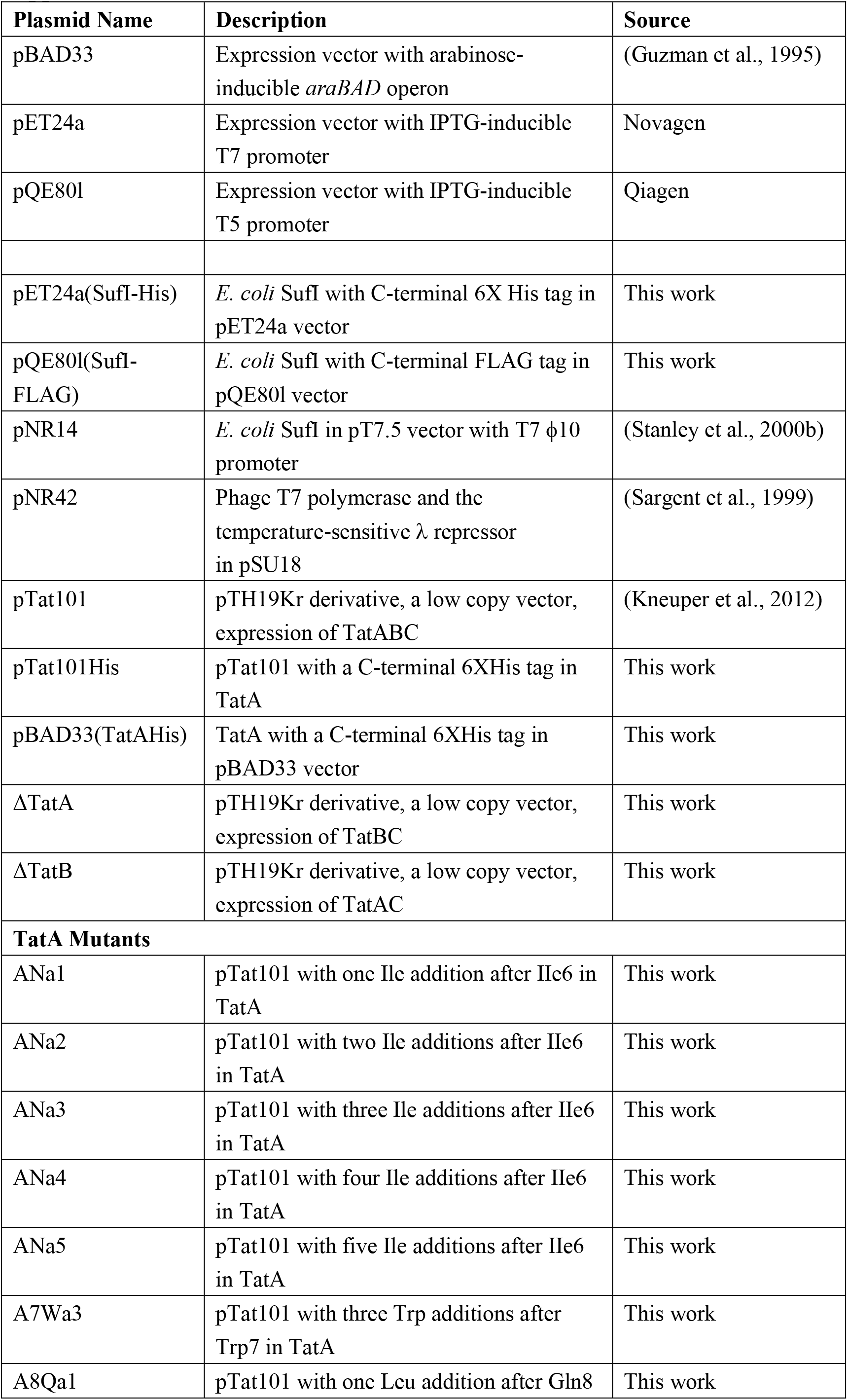
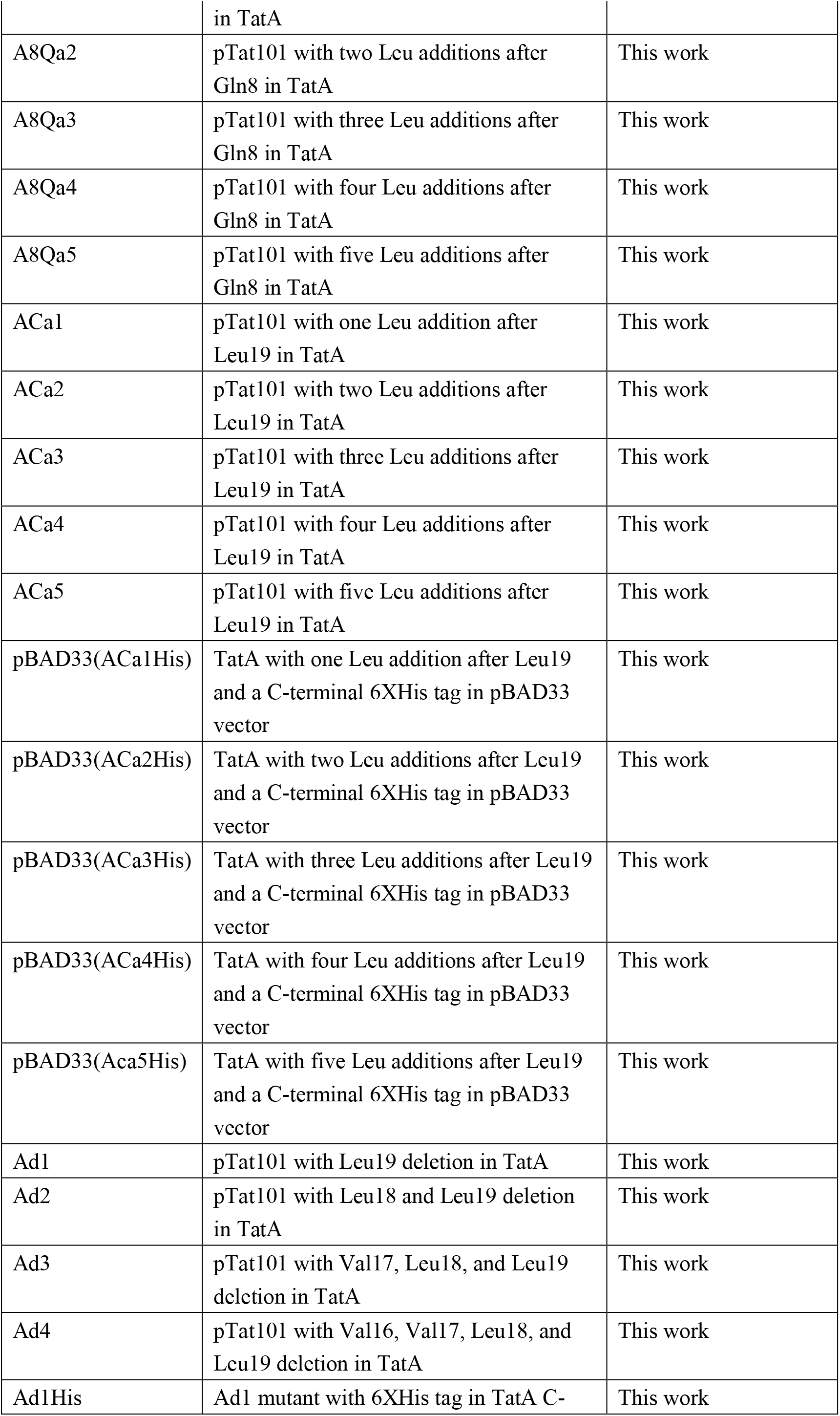
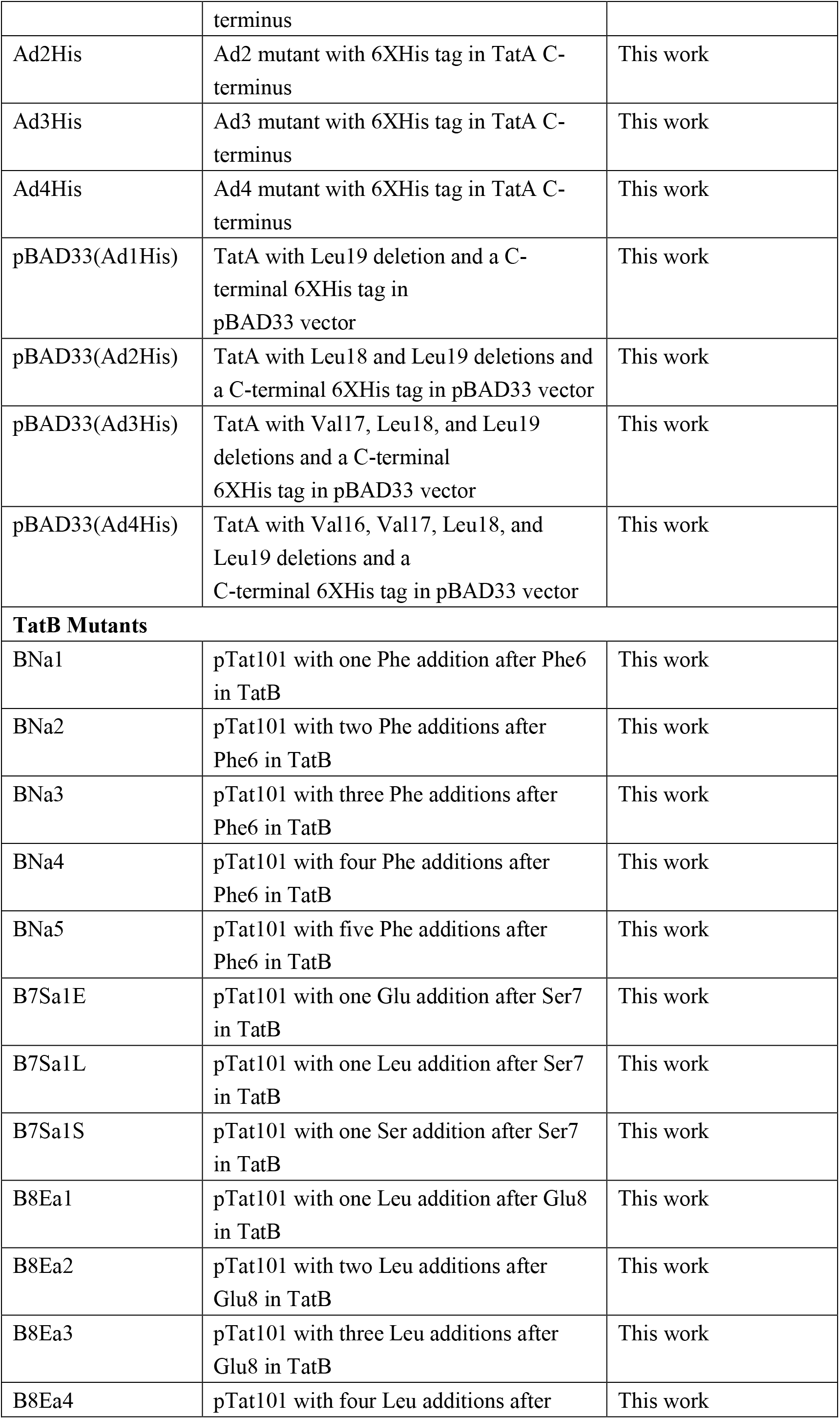
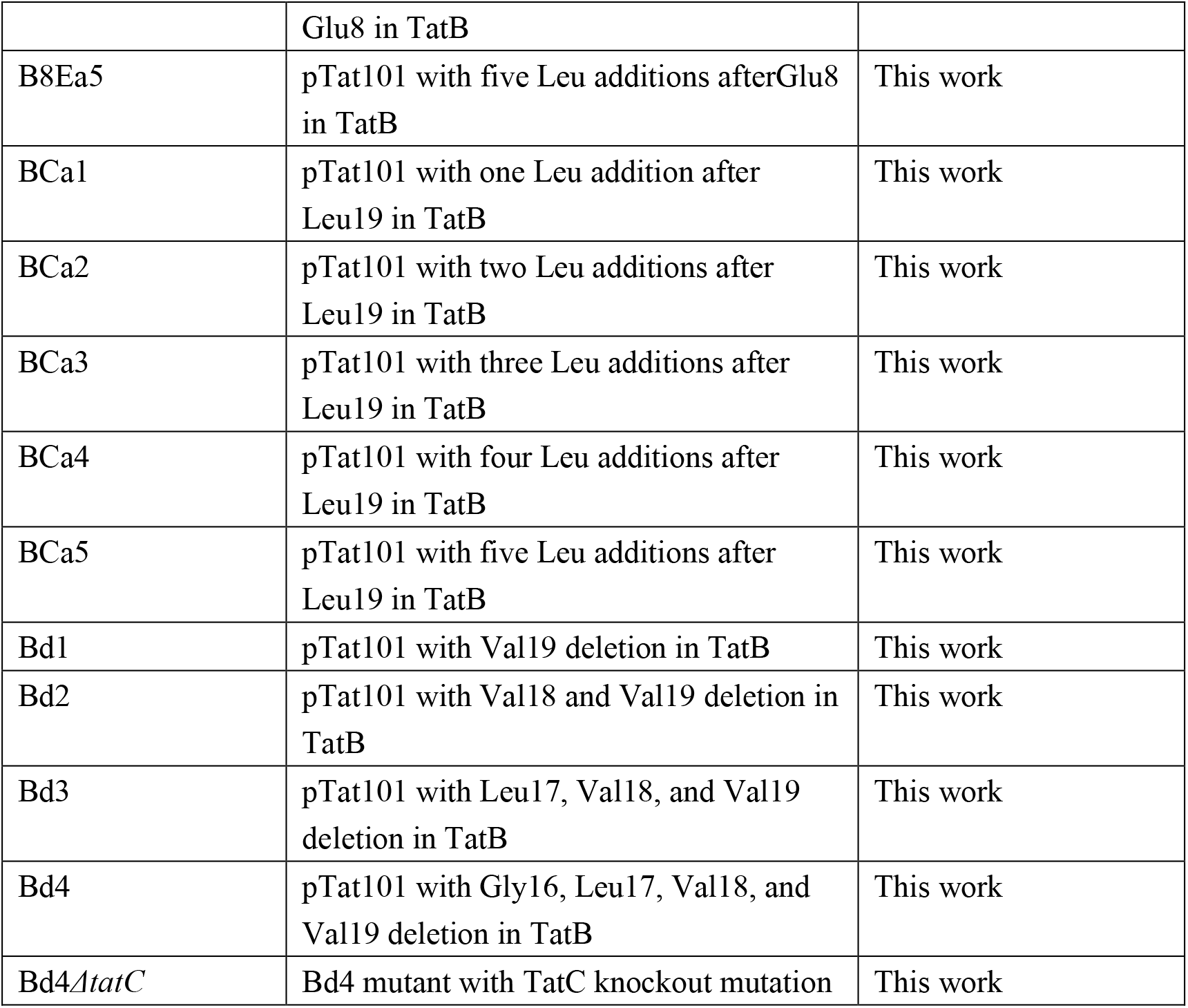
Plasmid information and mutants.

